# Nitrite secretion by cyanobacteria is controlled by the small protein NirP1

**DOI:** 10.1101/2023.08.04.552085

**Authors:** Alexander Kraus, Philipp Spät, Stefan Timm, Amy Wilson, Rhena Schumann, Martin Hagemann, Boris Macek, Wolfgang R. Hess

## Abstract

When the supply of inorganic carbon is limiting, photosynthetic cyanobacteria excrete nitrite, a toxic intermediate in the ammonia assimilation pathway from nitrate. While it has been hypothesized that the secreted nitrite represents excess nitrogen that could not be further assimilated due to the missing carbon, the underlying molecular mechanism has remained enigmatic. Here, we characterized a hitherto unannotated gene with homologs in the genomes of 485 different cyanobacteria that is upregulated under low carbon conditions and controlled by the transcription factor NtcA, a central regulator of nitrogen homeostasis. To understand its function, we ectopically overexpressed it in *Synechocystis* sp. PCC 6803, which resulted in a chlorotic phenotype, delayed growth, severe changes in amino acid pools, and nitrite excretion. Coimmunoprecipitation experiments revealed that this protein targets nitrite reductase, a central enzyme in the assimilation of ammonia from nitrate/nitrite, and was re-named to nitrite reductase regulator protein 1 (NirP1). Our results reveal that NirP1 is widely conserved in cyanobacteria and plays a crucial role in the coordination of C/N primary metabolism by targeting one of the central enzymes. In natural environments, the excreted nitrite will be utilized by other microorganisms; therefore, NirP1 ultimately impacts the activities and composition of the surrounding microbiome.

## INTRODUCTION

### Cyanobacterial Control of Nitrogen and Carbon Homeostasis

Some cyanobacteria, such as marine *Prochlorococcus* and *Synechococcus*, are of paramount importance as primary producers and for the global biogeochemical carbon (C) and nitrogen (N) cycles [1–3]. Other cyanobacteria, such as *Synechococcus* sp. PCC 7942 (*Synechococcus* 7942) or *Synechocystis* sp. PCC 6803 (*Synechocystis* 6803), have become model species for the CO_2_-neutral production of chemical feedstock and biotechnological processes driven by photosynthesis [4, 5]. The two most relevant nutrients for cyanobacteria are inorganic carbon (C_i_) and N. Most cyanobacteria assimilate combined inorganic N forms, such as ammonia (NH_4_), nitrate (NO_3_), nitrite (NO_2_) and urea; in addition, diazotrophic species can utilize N_2_ gas.

Surprisingly, several different cyanobacteria excrete nitrite under photoautotrophic growth conditions. Certain isolates of the marine cyanobacterium *Prochlorococcus* were reported to release up to 30% of their nitrogen uptake as extracellular nitrite when grown on nitrate [6]. The released nitrite is cross-fed by other microbes that cannot consume nitrate or produce nitrite. Moreover, it was demonstrated that cooccurring microbes could be beneficial for *Prochlorococcus* [7, 8]. In *Synechococcus* 7942, as well, approximately 30% of the produced nitrite were excreted in shifts to low carbon (LC) conditions, while no nitrite was excreted under high carbon (HC) conditions [9]. The coordination of C and N metabolism is regulated by several transcription factors and other regulatory proteins. An acute N scarcity is sensed by two central regulators of nitrogen assimilation in cyanobacteria, P_II_ and NtcA, by binding the key metabolite 2-oxoglutarate (2-OG) [10–14]. 2-OG connects C and N metabolism because it represents the main precursor for ammonia assimilation through the glutamine synthetase/glutamine oxoglutarate aminotransferase (GS/GOGAT) cycle in cyanobacteria and plants. Initially, GS incorporates ammonia into glutamate, thereby producing glutamine. The amino group from glutamine is then transferred onto 2-OG by GOGAT, yielding two glutamate molecules. As a consequence, the intracellular level of 2-OG starts to increase once these reactions slow down due to insufficient nitrogen supply; thus, 2-OG is an excellent indicator of the nitrogen status [15]. Binding of 2-OG stimulates the activity of NtcA, the main transcriptional regulator of genes for proteins involved in nitrogen assimilation [10, 11]. Depending on the 2-OG level, the P_II_ interaction protein X (PipX) switches from binding to P_II_ to interacting with NtcA, further enhancing the binding affinity of this complex to target promoters [12, 14]. In *Synechocystis* 6803, NtcA directly activates 51 genes, including transporters for nitrogen sources and GS/GOGAT, and represses 28 other genes after 4 h of nitrogen starvation [16].

In addition to transcription factors, multiple different proteins have been identified that interact with these factors or the involved enzymes and govern their functions directly. Archetypical examples for these proteins are the inhibitory factors IF7 and IF17, as their binding to GS leads to the inactivation of the enzyme when nitrogen supply exceeds demand [17]. The P_II_-interacting regulator of arginine synthesis (PirA) regulates the flux into the ornithine-ammonia cycle and therefore arginine synthesis [18, 19], while the P_II_-interacting regulator of carbon metabolism (PirC) functions as an inhibitor of 2,3-phosphoglycerate–independent phosphoglycerate mutase (PGAM), the enzyme that shifts newly fixed CO_2_ toward lower glycolysis [20]. PirC becomes sequestered by P_II_ at low 2-OG levels and released for PGAM inhibition at high 2-OG levels and therefore connects the control of carbon and nitrogen assimilatory pathways in a particular way [20].

We have previously analyzed the primary transcriptomes of *Synechocystis* 6803 and the closely related strain *Synechocystis* sp. PCC 6714 [21–23]. Based on these analyses, many more genes encoding unknown small, potentially regulatory proteins were computationally predicted. Experiments using a 3×FLAG epitope tag fused in-frame to the 3’ ends of the respective reading frames validated five of these to encode small proteins [24]. Subsequent experimental analyses using coimmunoprecipitation (co-IP) demonstrated that one of these small proteins, NblD, facilitates the degradation of phycobilisomes under nitrogen starvation conditions [25], while another one, called AtpΘ, is an inhibitor of the ATP synthase back reaction under low energy conditions [26]. Collectively, these findings suggest that cyanobacteria use a broad collection of small protein modulators to regulate metabolism according to external clues.

Here, we report a small, regulatory protein involved in the control of nitrogen assimilation in cyanobacteria in an unexpected way. This protein is 81 amino acids long in *Synechocystis* 6803, and its expression is upregulated following shifts from HC to LC conditions and downregulated upon nitrogen downshifts. Co-IP experiments identified the enzyme nitrite reductase (NiR) as the primary target of this protein. Following this lead further, we characterize the protein as a regulatory factor in the primary assimilation of nitrogen from nitrate/nitrite. Based on the observed phenotype, we named the regulatory protein “NirP1” for nitrite reductase regulator protein 1.

## RESULTS

### The ncr1071 Transcript Exhibits Coding Potential

The NtcA regulon in *Synechocystis* 6803 was reported to comprise 79 genes, including eight non-coding RNAs (ncRNAs) [16]. One of these ncRNAs, with the transcript ID ncr1071, was strongly downregulated upon N step-down [16]. This transcript extends from position 2215954 to 2217039 on the forward strand of the *Synechocystis* 6803 chromosome (GenBank accession no. NC_000911) and was assigned to the transcriptional unit (TU) 2296 [21]. When 10 different cultivation conditions were compared, TU2296 was found to be maximally transcribed under LC, while it was strongly repressed under N starvation conditions [21]. TU2296 overlaps the coding sequence of the *sll1864* gene encoding a potential chloride channel protein on the reverse strand by more than 500 nt (**Fig. 1A**).

**Fig. 1.**
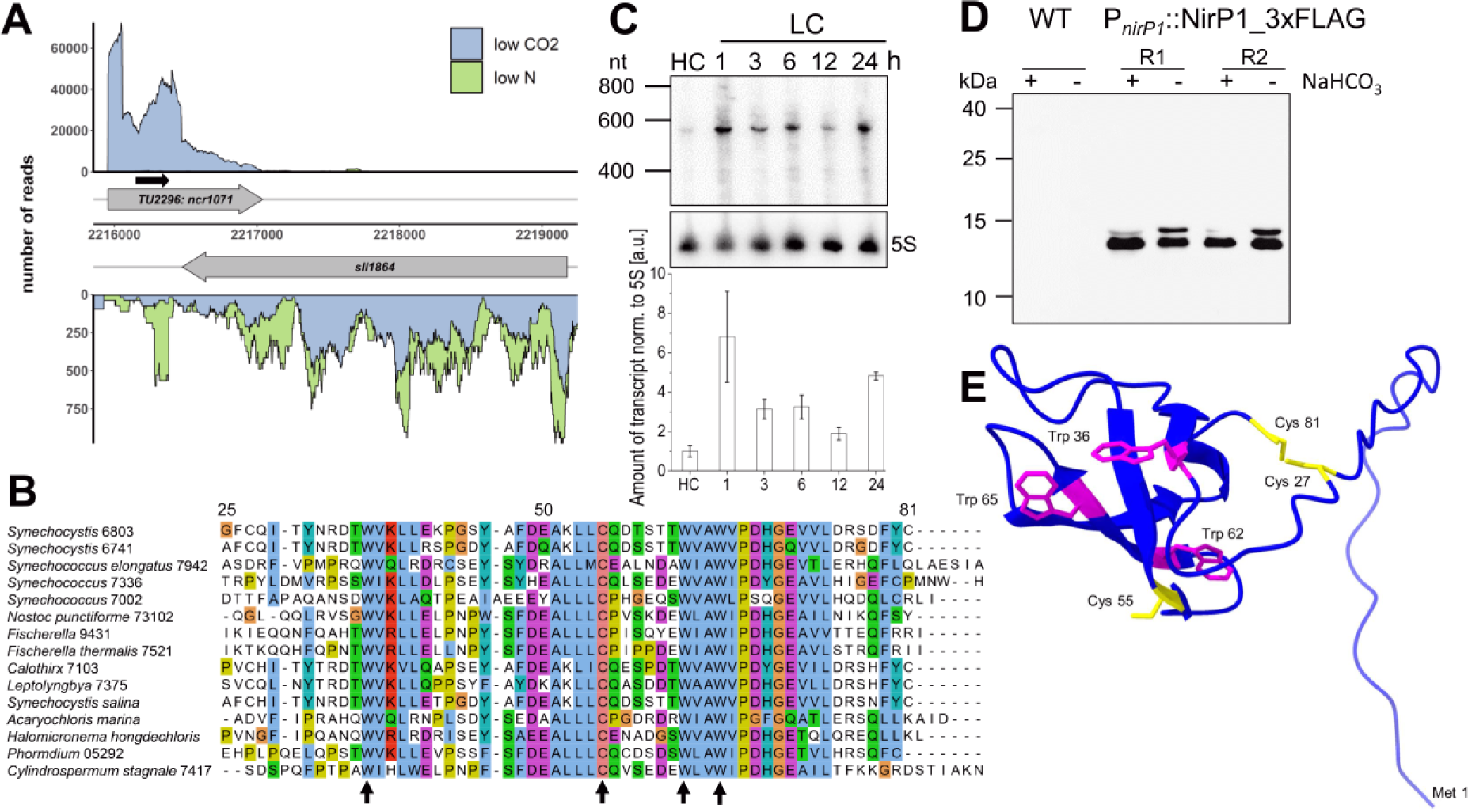
Genomic location and expression of *nirP1* in *Synechocystis* 6803. **A** The transcriptional unit (TU) TU2296 encompassing the coding sequence of *nirP1* (*ncr1071*) overlaps gene *sll1864* annotated as encoding a chloride ion channel protein. Plot at low CO_2_ (blue) versus nitrogen starvation (green). Transcriptional units (TUs) are indicated according to the previous annotation of the transcriptome and genome-wide mapping of transcriptional start sites [21]. The potential open reading frame of *ncr1071* coding for the NirP1 protein is indicated by a black arrow. **B** Alignment of selected potential NirP1 homologs from cyanobacteria belonging to four morphological subsections [28]. The last and most conserved part of *Synechocystis* 6803 NirP1 from position 25 to the end is shown. The cysteine and tryptophan residues conserved in all 485 potential homologs (**Supplemental Dataset 1**) are marked by arrows. The alignment was generated using ClustalW and visualized by Jalview. **C** Time course of *nirp1* mRNA accumulation in *Synechocystis* 6803 after cells precultivated in medium supplemented with 10 mM NaHCO_3_ (HC) for 3 hours were shifted to medium without a source of C_i_ (LC). The *nirp1* transcript was detected by a ^32^P-labeled single-stranded RNA probe. The membrane was rehybridized to a 5S rRNA probe as a loading control. The RiboRuler Low Range RNA ladder (Thermo Fisher Scientific) was used as the molecular mass standard. The relative amount of *nirp1* transcript normalized to the 5S rRNA is shown in the bar chart. The HC condition was set to 1, and all other signals were normalized to the HC condition. Two independent biological replicates were used. (*D*) NirP1 was detected in the presence or absence of C_i_ by Western blotting using anti-FLAG antiserum against tagged NirP1 under the control of its native promoter in two biological replicates (R1 and R2). The wild type was used as a control. Cultures were grown in medium supplemented with 10 mM NaHCO_3_ (+), washed and cultivated in carbonate-free medium (-) for 24 h. Prestained PageRuler^TM^ (Thermo Fisher Scientific) was used as a molecular mass marker. (*E*) NirP1 structure predicted by AlphaFold [31, 32] for the complete *Synechocystis* NirP1 protein, indicating the presence of four beta folds in the most conserved part. The four totally conserved amino acids as well as Cys27 and Cys81 are highlighted, tryptophan is shown in magenta and cysteine in yellow. The Met1 residue is also indicated for orientation.

Upon closer inspection, we discovered that the first segment of this transcript exhibits potential to encode an 81-residue protein with a calculated IEP of 4.59 and a molecular mass of 9.18 kDa. Evidence for the possible existence of this protein was provided independently by mass spectrometry data from a deep proteogenomics dataset [27]. Based on the results below, we named this protein “NirP1” for nitrite reductase regulator protein 1. Database searches identified 485 potential homologs of *nirP1* (**Supplemental Dataset 1**) in species belonging to all morphological subsections of cyanobacteria [28], including the well-studied model *Synechococcus* 7942, heterocyst-forming diazotrophic species and Section V strains, such as *Fischerella*. Homologous genes were detected in marine species with deviating pigmentation, such as *Acaryochloris marina* and *Halomicronema hongdechloris*, but not in *Prochlorococcus* and not in the endosymbiont chromatophore genomes of photosynthetic *Paulinella* species or the UCYN-A “Candidatus Atelocyanobacterium thalassa” endosymbionts, which can perform N_2_ fixation but lack photosystem II and phycobilisomes [29, 30]. Homologs were also not detected in the genomes of diatom-associated symbionts, *Calothrix rhizosoleniae* and *Richelia intracellularis*. These data suggest that homologs of *nirP1* were advantageous but not essential for free-living cyanobacteria with very different physiologies.

The vast majority of the 485 homologs identified in this work (**Supplemental Dataset 1**) are between 76 and 86 aa in length. The alignment of selected NirP1 homologs highlights the conservation in the last ∼50 aa and the presence of a centrally located, absolutely conserved single cysteine and three tryptophan residues (**Fig. 1B**).

To verify the inducibility of *nirP1* transcription by shifts to LC conditions, cultures of *Synechocystis* 6803 precultivated in medium supplemented with 10 mM NaHCO_3_ (high carbon, HC) were centrifuged and resuspended in medium lacking any source of C_i_. In Northern analysis, samples taken over 24 h showed a single signal of approximately 580 nt (**Fig. 1*C***) matching the length from the transcription start site (TSS) to the steep decline in the number of reads approximately 80 nt after the *nirP1* stop codon (**Fig. 1A**). The strongest signal was obtained 1 h after the shift with somewhat declining intensity at the later time points. At a weaker intensity, the *nirP1* mRNA was also detectable in the sample from high C_i_ (HC)-grown cells.

To directly validate the presence of NirP1, strains expressing NirP1 -3×FLAG fusions were constructed (**Supplementary Fig. S1**). Protein extracts were prepared and analyzed by SDS–polyacrylamide gel electrophoresis (SDS‒PAGE) and Western blotting using anti-FLAG antiserum. The calculated molecular mass of ∼12 kDa for the single detected signal was consistent with the sum of the calculated molecular masses of 9.18 for NirP1 and 2.86 for the 3xFLAG tag (**Fig. 1D**). No FLAG-tagged NirP1 was detected in the wild-type strain used as a control. From these experiments, we conclude that TU2296 encodes the small protein NirP1 and that *nirP1* transcription responds positively to shifts in C_i_ supply.

To generate a model for the NirP1 structure, we used AlphaFold [31, 32], which modeled NirP1 as a monomer and predicted four beta folds in the most conserved part of the protein. No structure was predicted for the N-terminal section of the protein, consistent with a lack of sequence conservation (**Fig. 1B**). Two cysteine residues (Cys27 and Cys81) in *Synechocystis* 6803 NirP1 were predicted to be in close proximity to each other, potentially forming a disulfide bond (**Fig. 1E**), whereas a third, highly conserved Cys55 was found between two beta strands in the folded part, exposed to the outside. However, no functional domains could be predicted based on the primary sequence or modeled structure.

### Two Transcription Factors are Involved in the Regulation of *nirP1* Expression

Based on literature reports [16], we speculated that *nirP1* expression could be negatively regulated by NtcA binding to a conserved binding site over the TSS. Moreover, we observed a tandem repeat element located further upstream consisting of two octameric repeats TTTGT(T/C)AA separated by a dinucleotide. To investigate the possible regulation through these sequence elements, the promoter of *nirP1* (P*_nirP1_*; from −70 to +30, TSS at +1) was engineered in a transcriptional fusion to drive the *luxAB* reporter genes. We used the native P*_nirP1_* sequence or a promoter with mutated, likely relevant nucleotides in the NtcA binding site (P_NtcA-Mut_) and in the upstream promoter element (P_Repeat-Mut_; **Fig. 2A**).

**Fig. 2.**
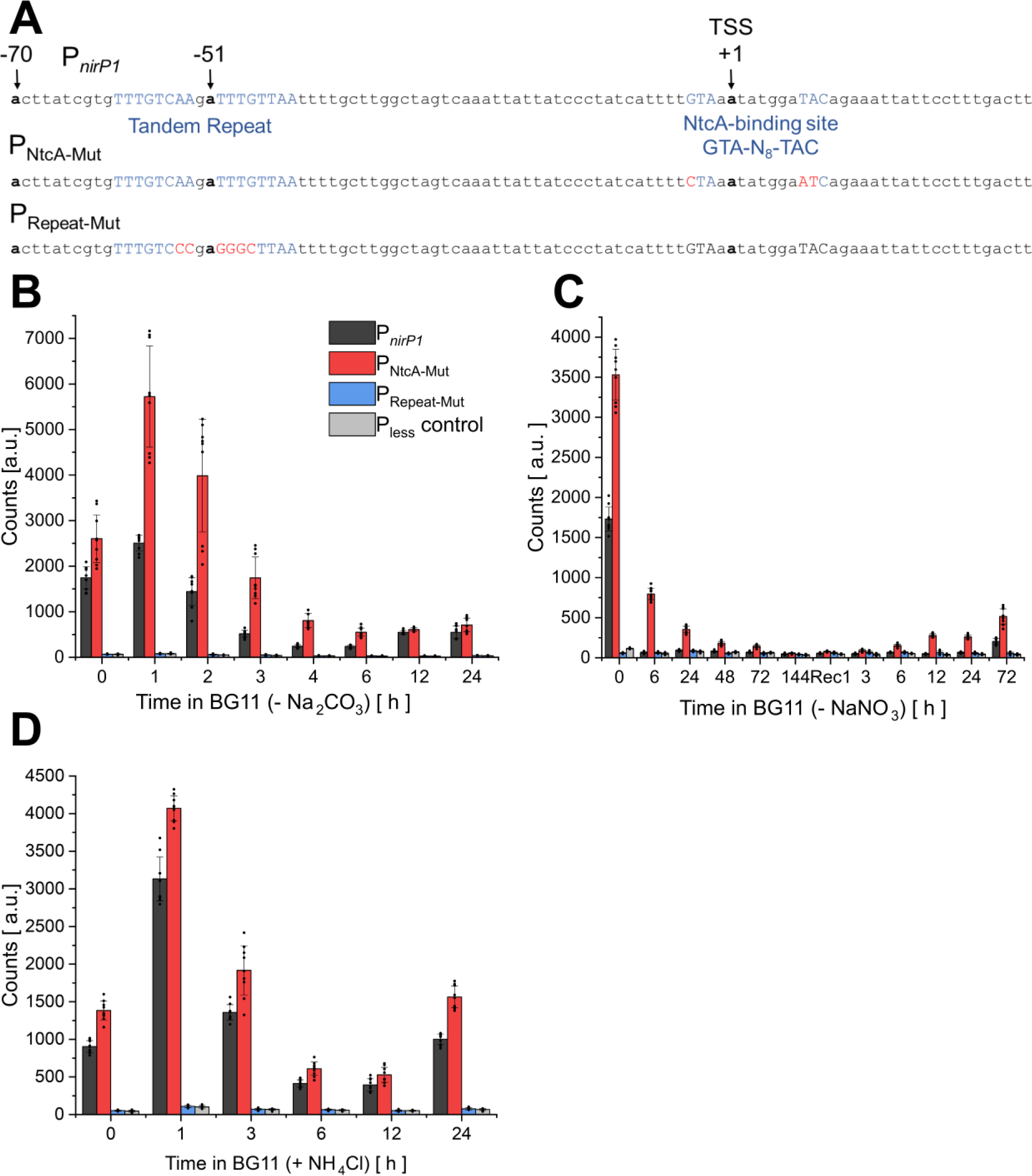
NirP1 expression is mediated through a C_i_-sensitive promoter and the transcription factor NtcA. **A** Top: Native P*_nirP1_* promoter sequence from position −70 to +30 (TSS at +1) used in a bioluminescence reporter strain harboring a P*_nirP1_*-*luxAB* transcriptional fusion. Functional promoter elements are colored blue. The previously determined [21] transcription start site (TSS) is indicated. Middle row: Mutated nucleotides in the NtcA binding site yielding promoter P_NtcA-Mut_ are colored red. Bottom: Mutated nucleotides in the repeat motif yielding promoter P_Repeat-Mut_ are colored red. Both mutated promoters were fused to *luxAB* and introduced into a neutral site in parallel with the P*_nirP1_*-*luxAB* construct. **B** Bioluminescence of the *Synechocystis* 6803 P*_nirP1_*-*luxAB*, P_NtcA-Mut_-*luxAB* reporter and P_Repeat-Mut_-*luxAB* strains and a promoter-less (P_less_) negative control. Cells were grown under HC conditions (BG11 supplemented with 10 mM NaHCO_3_) for 2 h and transferred to BG11 medium without a CO_2_ source to induce LC conditions for 24 h. (*C*) The strains were cultivated in BG11 (-N), and the chlorotic cultures were transferred back to standard BG11 (17.6 mM NaNO_3_) medium to start the recovery process seven days after nitrogen starvation was initiated. (*D*) Strains were cultivated in standard BG11 and then 10 mM NH_4_Cl was added for 24 h. Bioluminescence data are presented as the means ± SDs of n independent measurements including three biological replicates (= mean of values of independent transformants shown as dots, n = 3). Significance was calculated with a two-sample *t* test with unequal variance (Welch’s *t* test; *, P < 0.05; **, P < 0.01; ***, P < 0.001) between the strains at corresponding time points (details of statistical analysis **Supplementary Fig. S2** and **in Supplemental Dataset 2**).

In strains P*_nirP1_* and P_NtcA-Mut_, transcription was activated 1 h after a shift to medium lacking C_i_, followed by a decrease (**Fig. 2B, Supplementary Fig. S2**). This activation was significantly higher in the P_NtcA-Mut_ strains with the mutated NtcA-binding site. This effect was even more pronounced after a shift to nitrogen starvation conditions (**Fig. 2*C***). These results indicated the repressor function of NtcA for *nirP1*, consistent with previous ChIP-seq [16] and transcriptome data [21]. However, repression was still observed 6 h after the shift to nitrogen starvation, even when the binding motif was mutated rendering NtcA binding impossible, indicating the presence of additional regulatory mechanisms. It is possible that a second transcription factor is involved in the repression of *nirP1* during nitrogen starvation.

If the tandem repeat was mutated (P_Repeat-Mut_), a signal close to baseline was detected in all tested conditions (**Fig. 2B, C and D, Supplementary Fig. S2**). We conclude that this motif is essential for the transcription of *nirP1*, consistent with its position centered at -51.5 from the TSS at +1 (**Fig. 2A**), a region where frequently transcription factors bind to activate transcription.

Finally, transcriptional regulation was tested in response to the addition of ammonium (**Fig. 2D**). The signal increased ∼threefold after 1 h and then decreased, similar to the results obtained after the shift in LC conditions (**Fig. 2B**). These data agree with the results of a recent transcriptome analysis in which the *ncr1071* transcript was upregulated ∼threefold during acclimation to ammonium stress [33], indicating that the C/N balance is pivotal for the transcription of *nirP1*.

Collectively, these results demonstrated the complex transcriptional regulation of *nirP1*. We validated the transcriptional regulation of *nirP1* by NtcA and obtained new insight into the regulation of *nirP1* in shifts to nitrogen deprivation and with different C/N ratios. Due to the observed strong regulation, we hypothesized that NirP1 might be a regulator and is likely involved in the acclimation to changing carbon and nitrogen supply.

### Deletion and Ectopic Expression of NirP1 Leads to Phenotypical Changes

To test the potential functional importance of NirP1, we constructed the following strains: a gene deletion strain Δ*nirP1*; two overexpression strains in which the coding sequence of *nirP1* was fused to a C-terminal 3×FLAG tag under control of the copper-inducible P*_petE_* promoter on a plasmid pVZ322 introduced in a WT background (strain NirP1oex) and in the Δ*nirP1* background (strain Δ*nirP1*::oex); and a complementation strain with the same plasmid as in the overexpression strains, except that *nirP1* was controlled by its native promoter (Δ*nirP1*::*nirP1*).

Under nutrient-replete conditions, the Δ*nirP1* deletion mutant showed linear growth similar to the wild type. In contrast, both strains expressing *nirP1* under control of the P*_petE_* promoter showed severely reduced growth in liquid medium, irrespective of whether the plasmid was in the wild type or the Δ*nirP1* background (**Fig. 3A**). The reduced growth observed at high *nirP1* expression compared to wild type and Δ*nirP1* was also visible under LC, while the strain grew normally if *nirP1* was controlled by its native P*_nirP1_*promoter (**Fig. 3B**). Therefore, we concluded that the high expression of *nirP1* from the P*_petE_*promoter led to reduced growth. In addition, we noticed a pigmentation change, i.e., the cultures became chlorotic if *nirP1* was overexpressed from the P*_petE_* promoter (**Fig. 3C**). This effect was strictly dependent on the *nirP1* expression level because non-induction in the absence of added Cu_2_SO_4_ showed the same pigmentation as the wild type (**Supplementary Fig. S3A**). This phenotype was verified in room temperature absorption spectra. Compared to the spectra of the wild type and deletion mutant, the phycobilisome peak at approximately 625 nm as well as the chlorophyll peaks at 440 and 680 nm were decreased in the two strains that expressed *nirP1* from P*_petE_*(**Fig. 3D**).

**Fig. 3.**
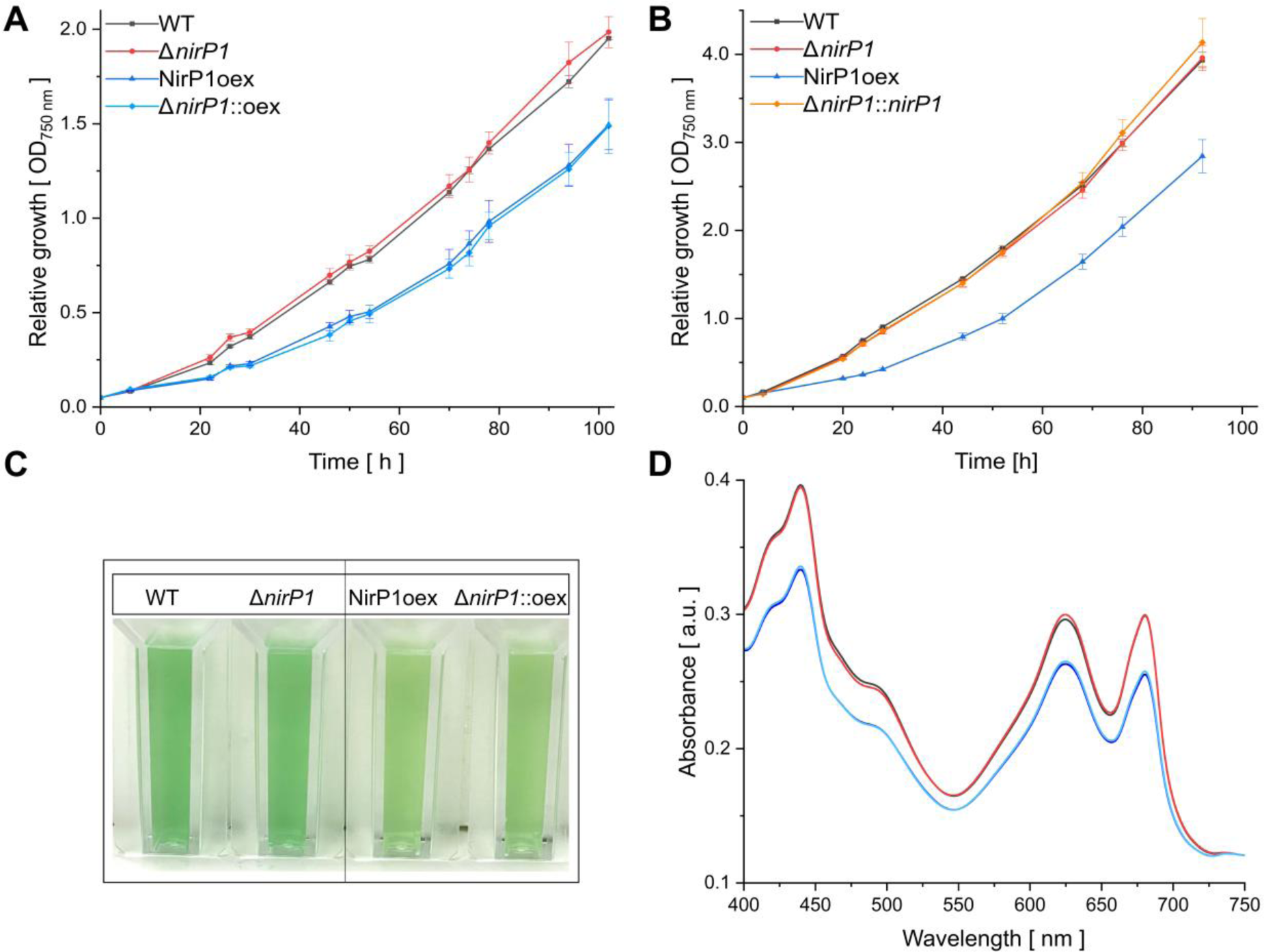
Phenotypical differences between *nirP1* mutant strains and the wild type in BG11 medium. **A** Growth of wild type (black), the Δ*nirP1* deletion mutant (red) and the strains expressing *nirP1* under control of the P*_petE_* promoter in the wild type background (dark blue, NirP1oex) and Δ*nirP1* background (light blue, Δ*nirP1*::oex). Data points represent the average of 3 biological replicates. **B** Growth in liquid medium after shifting to LC conditions. Strains were cultivated for 2 h in medium containing 10 mM NaHCO_3_, collected and transferred to medium without a CO_2_ source. The complementation strain (orange, Δ*nirP1*::*nirP1*) expressed *nirP1* under the control of its native *P_nirP1_* promoter in the Δ*nirP1* background. Other strains as in Panel (A); n=3. **C** Pigmentation phenotype of wild type and *nirP1* overexpressors in the presence of Cu_2_SO_4_ expressing *nirP1* for 24 h. All cultures were set to an OD of ∼0.8. **D** Room temperature absorption spectra for wild-type and *nirP1* mutants expressing *nirP1* for 24 h from the P*_petE_* promoter in the presence of Cu_2_SO_4_, normalized to wild-type OD_750_. Same strains and colors as in Panel (A). Spectra were recorded from three biological replicates and averaged.

To extend these results, drop dilution assays were performed to compare growth on plates. Again, the strain that overexpressed *nirP1* from the P*_petE_* promoter showed reduced vitality compared to the wild type and Δ*nirP1*, irrespective of the addition of 10 mM NaHCO_3_ (**Supplementary Fig. S3B**).

To summarize, the phenotypical effects due to ectopic NirP1 expression suggested that the amount of NirP1 played a critical role in the cell, while its absence did not lead to major phenotypic effects under the tested conditions.

### Metabolic Changes in the GS/GOGAT and Ammonia-Ornithine Cycle

Upon *nirP1* overexpression, the chlorotic phenotype (**Fig. 3C, D**) resembled the pigmentation changes characteristic of nitrogen-starved cells. Together with the observed expression control by NtcA (**Fig. 2**), this suggested a primary relationship between NirP1 and processes related to the assimilation of nitrogen, albeit the regulatory effect of LC.

Therefore, we tested whether the ectopic expression of NirP1 influences amino acid levels in the cell. Metabolites and amino acids were extracted in a time course experiment from wild type, Δ*nirP1*, and NirP1oex following a shift to LC. The total amount of soluble amino acids was significantly higher in NirP1oex at 12 and 24 h after transfer to LC than in the other two strains (**Fig. 4A**, full data in **Supplemental Dataset 4**). However, a closer look at the data revealed that the higher amino acid levels in NirP1oex were mainly caused by the accumulation of a single amino acid, glutamate (**Fig. 4B**). The glutamate level in *Synechocystis* 6803 is normally approximately 10 times higher than that of any other amino acid. Therefore, the increased accumulation of glutamate alone was responsible for the measured higher total amino acid amount in NirP1oex. However, for the direct product of glutamate amination by GOGAT, glutamine, we measured lower levels in NirP1oex at 3 and 24 h after the shift to LC (**Fig. 4B**). This result is consistent with previous analyses in which a decrease in glutamine content was observed in LC shifts (52). In addition, we found that the concentrations of citrulline and arginine, which are produced from glutamine via the ornithine-ammonium cycle (OAC), were lowered in NirP1oex at 3, 12 and 24 h after the induction of expression (**Fig. 4C**). In contrast, the aspartate levels were increased, consistent with its synthesis from glutamate by glutamate aspartate aminotransferase. We conclude that the upregulation of NirP1 caused several changes in amino acid levels related to the synchronization between C and N metabolism. As the OAC is involved in nitrogen storage and remobilization [36], these results support that NirP1 plays a role as a regulator in nitrogen metabolism.

**Fig. 4.**
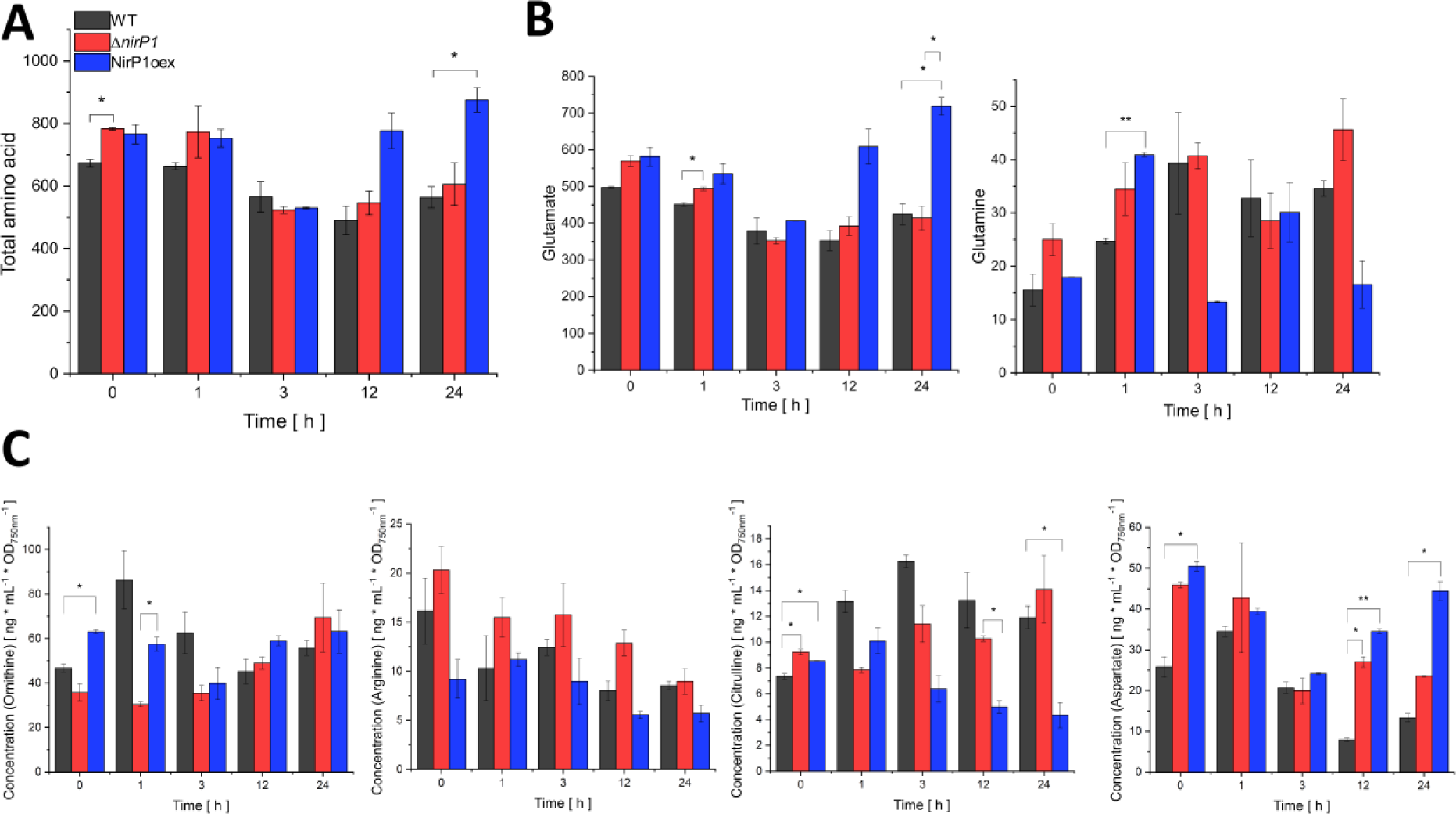
Differences in the accumulation of key intermediates in C/N primary metabolism due to the availability of NirP1 during shifts to LC. A Total amino acid content. B Glutamate and glutamine concentrations. C Concentrations of ornithine-ammonium cycle intermediates and of aspartate. Metabolite content was measured for Δ*nirP1*, NirP1oex and wild type (WT). The concentrations are given in mM for total amino acids, glutamate and glutamine, all other concentrations are given in ng * mL^-1^ * OD750 nm^-1^. Two biological replicates of all strains were used and averaged. Significance was calculated with the two-sample t test with unequal variance (Welch’s t test; *, P ≤ 0.05; **, P ≤ 0.01; ***, P ≤ 0.001) for each strain and between the strains at corresponding time points (details in Supplemental Dataset 4).

### NirP1 Interacts with Ferredoxin-Nitrite Reductase

Coimmunoprecipitation (co-IP) was successfully used in the past to identify the interacting partners of *Synechocystis* 6803 small proteins NblD, AtpΘ, and SliP4, thereby providing essential hints for their specific functions [25, 26, 37]. Therefore, we used a strain expressing epitope-tagged NirP1 to obtain insight into the molecular mechanism that involves NirP1. After 24 h of induction, total cell extracts of three replicates of NirP1oex were analyzed by SDS‒PAGE and Western blotting using anti-FLAG antisera. The size of the NirP1-3xFLAG signal in the Western blot matched the sum of the calculated molecular masses of 9.81 kDa for NirP1 and 2.86 for the 3xFLAG tag. No signal was detected in the wild type used as a negative control (**Fig. 5A**).

**Fig. 5.**
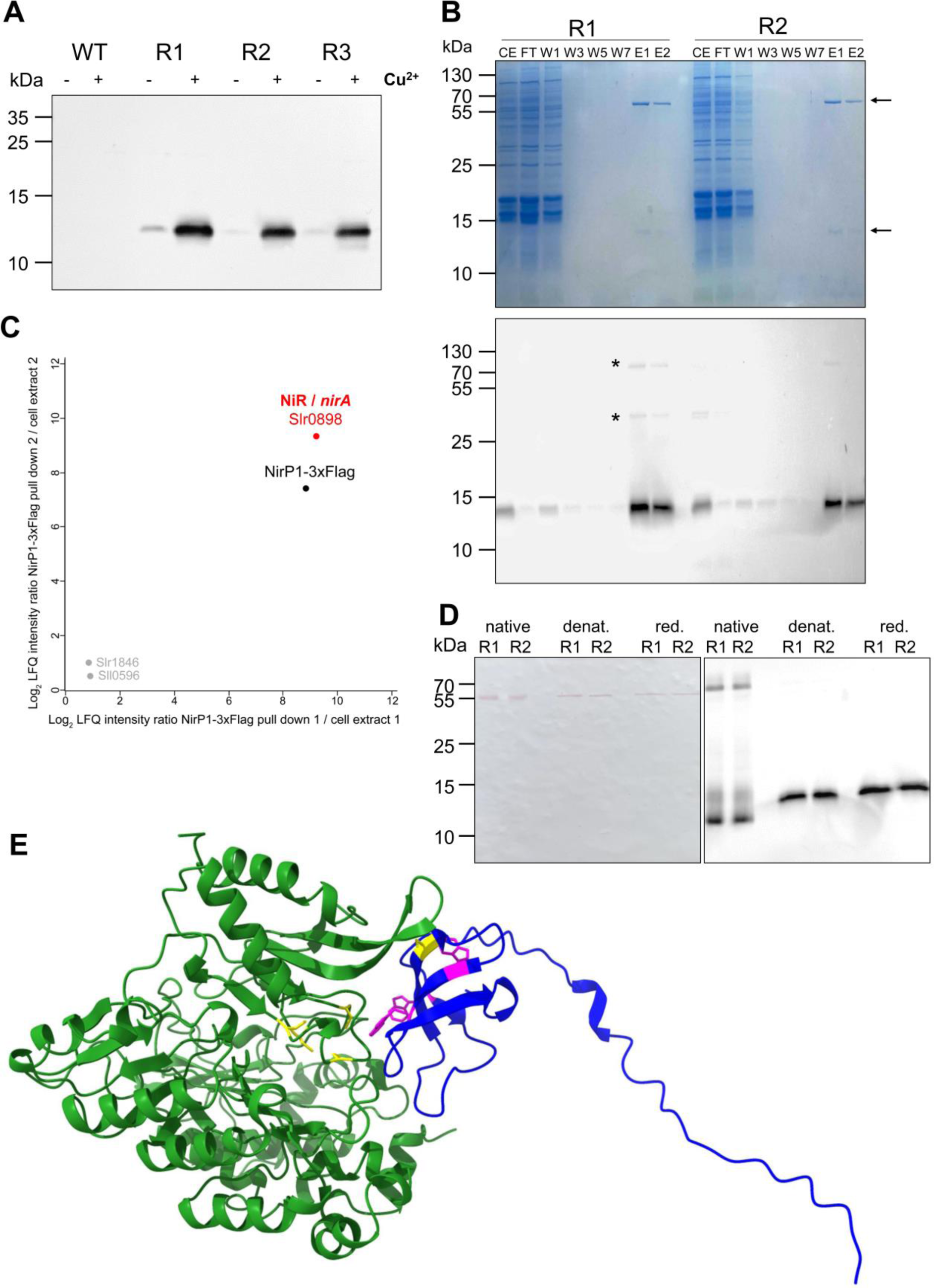
NirP1 expression and pull-down analysis. **A** Detection of NirP1 after 24 h of induction with 2 µM Cu_2_SO_4_ by Western blotting using anti-FLAG antiserum against tagged NirP1 in three biological replicates (R1 to R3). The wild type was used as a control. **B** NirP1 pull-down analysis of two biological, independent replicates (R1 and R2). Upper panel: Coomassie-stained SDS gel, the NirP1 signal and a band indicating a prominent coeluting protein of ∼60 kDa are marked with arrows. Fractions are numbered and labeled CE, cell extract; FT, flow-through; W, wash; E, elution. Lower panel: Immunological identification of the lower band as NirP1. **C** Mass spectrometry-based analysis of NirP1 co-IP. Scatter plot of log_2_-transformed LFQ protein ratios (NirP1-3xFlag co-IP/cell extract) from two independent replicates. Displayed are proteins with a higher abundance in NirP1-3xFlag co-IP elution fractions compared to the initial cell extract. The main interacting protein was identified as ferredoxin-nitrite reductase (NiR), which is encoded by the *nirA/slr0898 gene*. **D** Western blot of NirP1 co-IP fractions using native, denaturing or reducing conditions. All samples were loaded in biological replicates R1 and R2. (*E*) NirP1-NiR (WP_010873675.1) interaction predicted by AlphaFold [31, 32] using their full-length sequences. The interaction modeled for the NiR and NirP1 homologs from *Synechococcus* 7942 is shown in **Supplementary Fig. S5B.**

NirP1-3xFLAG and potential bound interaction partners were coimmunoprecipitated from the lysate using anti-FLAG resin (**Fig. 5B**). Elution fractions of the protein pull-down showed NirP1 in Coomassie-stained protein gels and an additional band of higher molecular mass in both replicates. Because signals were obtained in the Western blot at these two and a third position, this result pointed at potential interaction partners of NirP1 that were copurified with the bait protein. These additional proteins were still present after 8 washing steps, indicating a strong interaction with NirP1 (**Fig. 5B**).

Next, we subjected whole cell lysates and eluate fractions to liquid chromatography-tandem mass spectrometry (LC‒MS/MS)-based proteome analysis for the identification of coimmunoprecipitated proteins. In addition to NirP1, a single other protein was strongly enriched, which was identified as the 56 kDa ferredoxin-nitrite reductase (NiR), encoded by *nirA* (*slr0898*) (**Fig. 5C**). Overall, eight and 72 nonredundant peptides of the NirP1-3xFLAG protein and of NiR were detected, yielding high respective sequence coverages of 94% and 98% (**Supplementary Table S3, S4 and S5**). The combined molecular masses of NirP1 and NiR are compatible with the uppermost signal in the Western blot in **Fig. 5B**. These results indicated that NiR was a specifically enriched protein that coimmunoprecipitated with the NirP1-3xFLAG protein under the conditions used.

To validate the potential complex formation between NirP1 and NiR, Western blot experiments were performed under different reducing conditions. Under native conditions, two bands appeared. The signal at the lower molecular mass matched the predicted molecular mass of monomeric NirP1, while the upper signal was consistent with the sum of NirP1 and NiR of ∼70 kDa. After the samples were denatured by heating or reduced through an addition of dithiothreitol (DTT) and β-mercaptoethanol, the ∼70 kDa signal disappeared. The single remaining band migrated higher than the band of the native form, indicating that it was the monomeric and completely unfolded NirP1 protein (**Fig. 5D**).

Modeling by AlphaFold [31, 32] predicted NirP1 binding to the NiR substrate binding pocket next to its catalytic center with the iron-sulfur cluster where the reduction of nitrite takes place (**Fig. 5E**). Interestingly, none of the NirP1 cysteine residues, but two of the three totally conserved tryptophan residues (W36 and W65), were predicted close to the catalytic center. Thus, these amino acids potentially play a functional role.

### Secretion of Nitrite is Triggered by NirP1

To test if the action of NirP1 might lead to the excretion of nitrite in LC conditions, as previously reported for *Synechococcus* 7942 [9], we first verified if *Synechocystis* 6803 showed this effect as well. Indeed, after 2 h at LC, the *Synechocystis* 6803 wild type, but not the Δ*nirP1* deletion mutant, started to excrete nitrite into the medium (**Fig. 6A)**. After 24 h at LC, the concentration of nitrite in the medium of the wild-type strain was ∼5 µM. However, this concentration was with ∼75 µM much higher for the *nirP1*-overexpressing strain (**Fig. 6A, B**). All three results (Δ*nirP1,* wild type, NirP1oex) suggested that the excretion of nitrite into the supernatant was related to the presence of NirP1 in the cell. This effect was even more pronounced by the ectopic overexpression of NirP1 in BG11 medium aerated with 5% (v/v) CO_2_ leading to an ∼150-fold higher accumulation of nitrite in the supernatant of NirP1oex compared to wild type and Δ*nirP1* grown samples (HC in **Fig. 6B**). After the cells were washed and resuspended in fresh medium, the overexpressor immediately excreted nitrite again (**Fig. 6B**). These data establish that NirP1 triggers the secretion of nitrite.

**Fig. 6.**
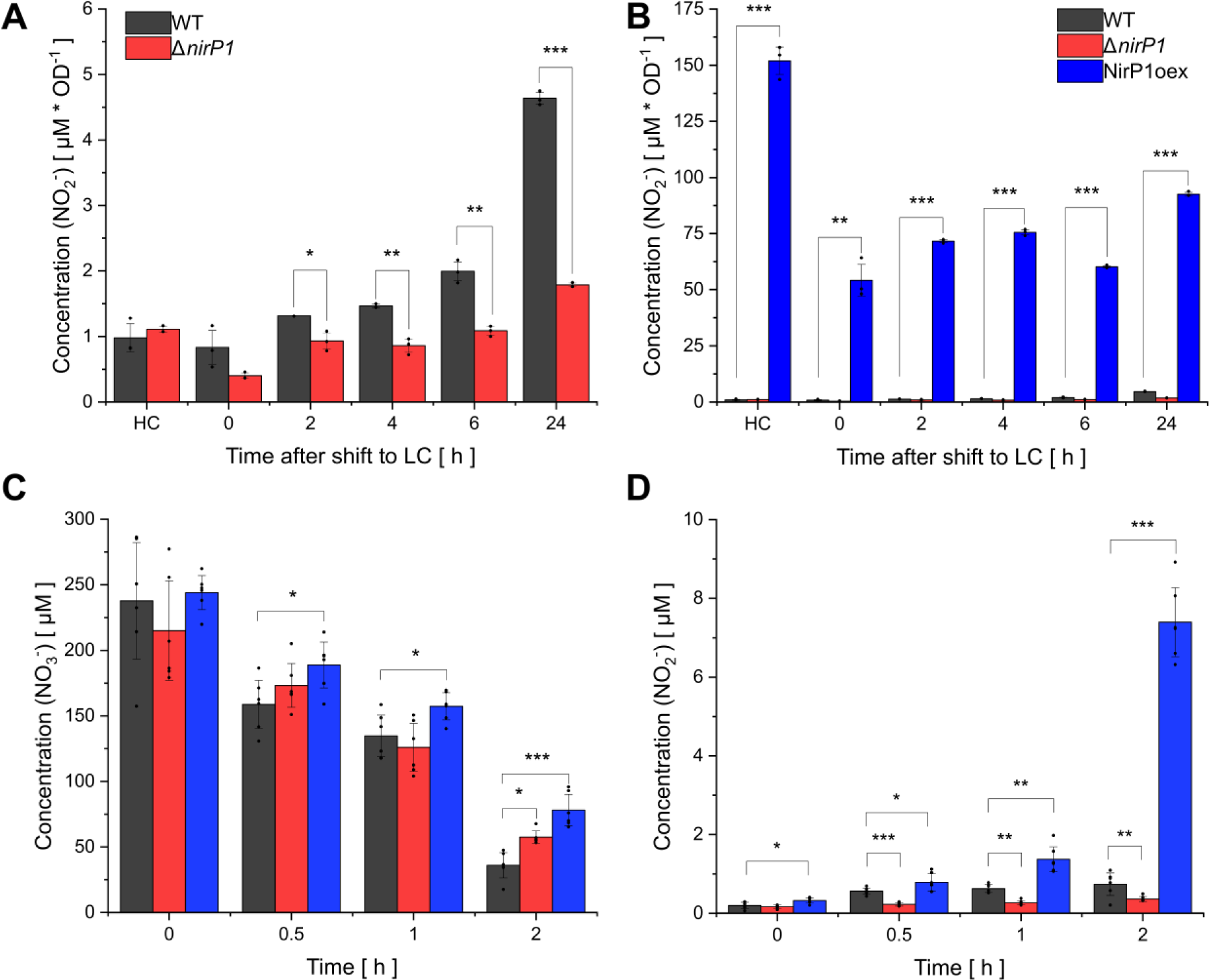
Secretion of nitrite is triggered in the presence of NirP1. **A** *Synechocystis* 6803 strains were grown photoautotrophically in BG11 medium with 5% (v/v) CO_2_ in air as high CO_2_ (HC), harvested and resuspended in new medium and aerated with 0.04% (v/v) CO_2_ in air to initiate LC conditions. The wild type started to excrete nitrite in LC conditions when *nirP1* was expressed (see Fig. 1C). **B** Same data as in Panel (A) but at different scales to include the results from the NirP1oex strain. Accumulation of nitrite in the supernatant was followed over time. Data are presented as the means ± SDs of biological triplicates. Significance was calculated using a two-sample *t* test with unequal variance (Welch’s *t* test; ns, no significance; *, P < 0.05; **, P < 0.01; ***, P < 0.001) between the strains at corresponding time points. **C** The consumption of nitrate, measured as the decrease in the concentration of nitrate in the supernatants after pelleting the cells by centrifugation. Strains were grown in BG11, washed with nitrogen-free BG11 (BG11 –N), set to OD = 1 and resuspended in BG11 containing 200 µM nitrate. **D** The production and secretion of nitrite into the supernatant measured in the same samples as in Panel (C) for nitrate. Full details are provided in **Supplemental Dataset 3.**

To ensure that the accumulated nitrite was a product of absorbed and reduced nitrate, the uptake of nitrate and subsequent secretion of nitrite were measured in parallel. The cultures were first grown in nitrogen-free BG11 medium and then 200 µM nitrate was added. In the control experiment, an additional 2 mM ammonia was added because nitrate uptake in *Synechocystis* is inhibited in the presence of ammonia and the transcription of nitrate and nitrite reductase genes is repressed. After 2 h, the overexpressor led to significant differences in nitrite concentration in the medium compared to wild type and Δ*nirP1*. In addition, the overexpression seemed to slow the uptake of nitrate (**Fig. 6C, D**). Here, the wild type, although less pronounced than the overexpressor, also excreted nitrite into the medium, as observed in the LC shift experiment before. In contrast, in the control experiment with additional ammonium, no nitrate uptake and no secretion of nitrite into the medium were observed (**Supplementary Fig. S4A, B**). These results demonstrated that the accumulation of nitrite in the medium of the wild type and overexpression strains resulted from nitrate uptake, the subsequent reduction to nitrite by NR and the inhibition of NiR by NirP1. Approximately 5% of assimilated nitrate was excreted as nitrite into the medium by *Synechocystis* in the first 2 h (**Fig**. **6C, D**). Compared to *Prochlorococcus* and *Synechococcus* 7942, which were reported to excrete up to 30% of the assimilated nitrate as nitrite [6, 9], the nitrite excretion of *Synechocystis* 6803 was less effective and the remaining nitrite was stored inside the cell. Surprisingly, the remaining nitrite did not seem to poison the cells.

## DISCUSSION

Nitrate is one of the most abundant nitrogen-containing nutrients for cyanobacteria, while the preferred source for nitrogen assimilation is ammonium. Therefore, complex regulatory mechanisms occur to regulate nitrogen assimilation and uptake, e.g., genes of nitrate assimilation are repressed if ammonium is available [38, 39]. When nitrate is the only nitrogen source available, it is taken up and reduced in two steps to nitrite and ammonium, which is then incorporated into glutamate in the GS/GOGAT cycle (**Fig. 7**). The reduction of nitrate to nitrite is catalyzed by the enzyme NaR and the subsequent reduction to ammonium by NiR. NirP1, a small regulatory protein, is upregulated in LC conditions in *Synechocystis* 6803 (**Fig. 1A**). Its elevated expression, as shown in this study, leads to the accumulation of nitrite in the medium. We identified NiR as the primary binding partner for NirP1 (**Fig. 5C**). Therefore, the results can be explained if the NirP1:NiR interaction directly leads to the accumulation of nitrite in the cell, which is toxic and therefore becomes rapidly exported (**Fig. 7**). Excess internal nitrite has been shown to inhibit photosynthesis, particularly the activity of photosystem II in *Synechocystis* 6803 [40]. Comparable responses were reported for nitrogen-starved *Synechocystis* 6803 mutants defective in glycogen synthesis that excreted 2-OG and pyruvate [41].

**Fig. 7.**
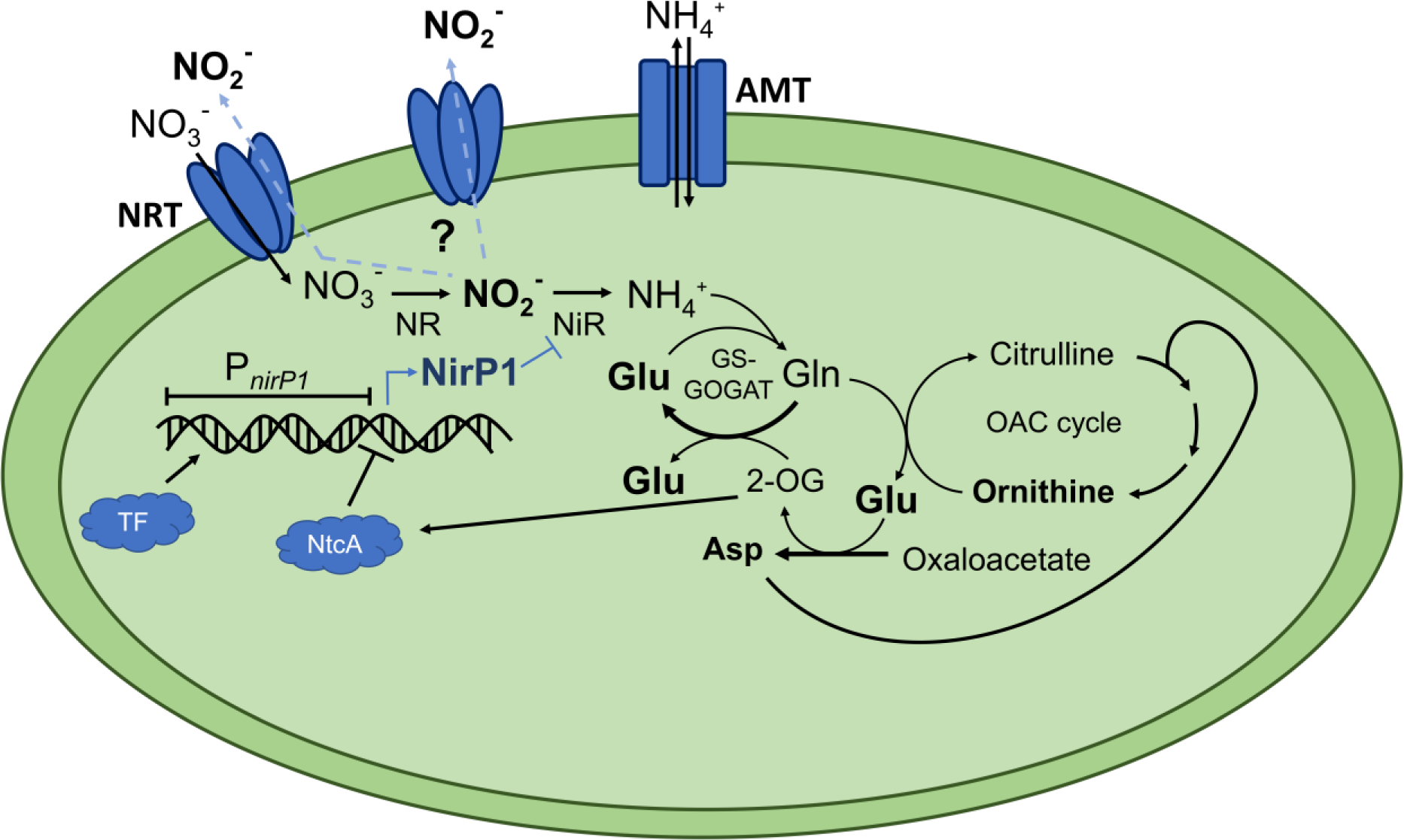
NirP1 is a regulator at a central position in the assimilation of inorganic nitrogen. Cyanobacteria can assimilate nitrogen from simple compounds, such as ammonium, nitrate, nitrite and urea. During nitrate assimilation, intracellular nitrate is reduced in two steps to nitrite and ammonium, which is then incorporated into glutamate in the GS/GOGAT cycle or into carbon skeletons during the synthesis of amino acids by transaminases. The reduction of nitrate to nitrite is catalyzed by the NR and the subsequent reduction to ammonium by the NiR. Transcription of the regulatory factor NirP1 is controlled by an unknown transcription factor and by NtcA. 2-OG is a corepressor of NtcA. Therefore, an increased 2-OG level leads to the repression of *nirP1* transcription, while a low level of 2-OG leads to its activation. NirP1 binds to the enzyme nitrite reductase (NiR), leading to the inhibition of nitrite reduction. The accumulating nitrite is then rapidly exported by an unidentified export system.

Several factors control primary nitrogen metabolism, such as the transcription factor NtcA or the P_II_ regulator and proteins interacting with them [10–14, 17–19]. However, unlike several other regulatory proteins that modulate the activity of other regulators (e.g., PirA and PipX), NirP1 targets one of the key enzymes of nitrogen assimilation metabolism directly. This is reminiscent of the direct inhibitory role of the small GS inhibitory factors IF7 and IF17 (27). These findings suggest that a higher number of relatively small proteins might exist that modulate or modify the activity of relevant enzymes and await characterization.

To examine the possible mode of action, we further predicted the interactions between NiR and other proteins. In *Synechocystis* 6803, ferredoxin Fd1 (*ssl0020, petF*) acts as an electron donor for the reduction of nitrite [42]. Therefore, we modeled the interaction of NiR with NirP1 (**Fig. 4E**) and Fd1 (**Supplementary Fig. S5A**). The results suggested that both proteins compete for the binding pocket of NiR. By binding NirP1 instead of Fd1, the binding pocket is blocked for electron transfer and the subsequent reduction of nitrite to ammonium. Thus, this could be a possible mechanism by which NirP1 inhibits NiR, leading to the secretion and accumulation of nitrite in the medium during NirP1 expression.

Transcription of *nirP1* is controlled by an unknown transcription factor and by NtcA. If 2-OG cannot be aminated efficiently in the GS-GOGAT cycle due to nitrogen starvation, carbon oversupply, or a combination of both, it will accumulate in the cell. The increased 2-OG level is sensed by PII and transferred to NtcA, leading to stronger promoter binding and the repression of *nirP1* transcription. Therefore, NirP1 is at the confluence of C and N assimilation and is effectively a regulator of the C/N balance. In addition to the NtcA binding site, we found another putative promoter element at position -43 to -60 (**Fig. 2A**), the mutation of which rendered the *nirP1* promoter inactive. Therefore, this element is likely recognized by a positively acting transcription factor, although its identity was not further investigated in this work and should be studied in the future.

It was previously reported that during photoautotrophic growth in shifts to LC conditions, *Synechococcus* 7942 excreted nitrite into the medium [9]. As the fundamental reason for this observation, a decline in the enzymatic activity of NiR was identified that started within 30 min. This decline was not a consequence of changed *nirA* expression because the observed decrease in NiR activity under LC conditions was substantially larger than that observed when the cultures were treated with rifampicin to inhibit transcription. Instead, a mechanism was postulated that directly inactivated NiR under LC conditions [9]. Our data were obtained in *Synechocystis* 6803, but a close NirP1 homolog is found in *Synechococcus* 7942 (**Fig. 1B**), and an almost identical interaction with NiR was predicted (**Supplementary Fig. S5B, C**). Therefore, we have likely identified the mechanism underlying previously observed NiR inhibition and nitrite excretion.

According to our data, NirP1 expression is upregulated under LC conditions and then participated in the inhibition of NiR, leading to nitrite excretion and accumulation. Surprisingly, the regulation occurs at an unexpected point (**Fig. 7**), as controlling N assimilation during nitrate uptake or during nitrate reduction seems more intuitive. Here, uncontrolled nitrate uptake led to light-dependent nitrite excretion under low light conditions [43]. Therefore, the availability of electrons could play a role because the reduction of nitrite requires six moles of reduced ferredoxin, while the reduction of nitrate requires only two. Another consideration is the possible beneficial effect of nitrite excretion on cooccurring microbes in the natural environment. Certain *Prochlorococcus* release up to 30% of their N uptake as extracellular nitrite when grown on nitrate [6]. *Prochlorococcus* typically contains the most compact genomes of any free-living cyanobacteria, which was linked to its quasi homeostatic environment [44–46]. Because we did not identify NirP1 homologs in *Prochlorococcus,* the underlying mechanism probably operates without NirP1. In other cyanobacteria, exposed to greater variations in the environment, the involvement of NirP1 has likely led to a more accurate regulation. In any case, the release of a N cycle intermediate, such as nitrite, should have tremendous relevance for certain other members of the microbiome. Our results show that homologs of NirP1 are widely distributed among cyanobacteria, expanding our understanding of how carbon and nitrogen metabolism become integrated in these organisms.

## MATERIALS AND METHODS

### Cultivation Conditions and Spectrometric Measurements

Strains were maintained BG11 or in copper-free BG11 [28] supplemented with 20 mM TES pH 7.5 under continuous white light of 50 µmol photons m^-2^ s^-1^ at 30 °C. Mutant strains containing pVZ322-derived plasmids were cultivated in the presence of 50 µg/mL kanamycin, while Δ*nirP1* was maintained at 20 µg/mL streptomycin. The cultures for overexpression of NirP1 using the P*_petE_* promoter for pull-down experiments and MS measurements were grown either in BG11 or under CellDeg® conditions supplemented with 2 µM CuSO_4_ as previously reported [47, 48]. Cultures in growth experiments and phenotypic assays were grown in BG11 medium supplemented with copper but without antibiotics to prevent any possible side effects.

Whole-cell absorption spectra were measured using a Specord® 250 Plus (Analytik Jena) spectrophotometer at room temperature and were normalized to 750 nm. Cultures were measured in triplicate. Cultures at an OD_750_ > 1 were diluted with 1× BG11 prior to taking the absorption spectra.

### Construction of Mutant and Overexpression Lines

*Synechocystis* 6803 PCC-M [49] was used as the wild type and background for the construction of mutants. Knockout mutants were generated by replacing the *nirP1* (*ncr1070*) coding sequence with a streptomycin resistance cassette (*aadA*) [50] by homologous recombination and using pUC19 as a vector for subcloning. The construct for gene replacement was generated by AQUA cloning [51]. Details of primer sequences and plasmids are provided in **Supplementary Tables S1 and S2**. Segregation of *nirP1* and Δ*nirP1/aadA* alleles was checked by colony PCR using the primers Seg_Norf9-ncr1070-fwd/rev (**Supplementary Fig. S1**).

To complement the knockout and create a NirP1-overexpressing strain, *nirP1* was inserted into the self-replicating pVZ322 plasmid under control of the Cu^2+^-inducible P*_petE_* promoter or under control of its native promoter (P*_nirP1_*). The inserts for the overexpression *of nirP1* were assembled into pVZ322 predigested by *Xba*I and *Pst*I for 16 h at 37 °C by AQUA cloning. The constructs were checked using the primers P_AK16 and P_AK17 (**Supplementary Fig. S1**). The sequence-verified plasmids were introduced into *Synechocystis* 6803 by electroporation [52]. The expression of *nirP1* was checked by Western hybridization using ANTI-FLAG antisera.

### Promoter Analysis

To verify *nirP1* promoter activity in vivo in *Synechocystis* 6803, the nucleotides −70 to +30 (TSS at +1) were fused to *luxAB* reporter genes and integrated into the genome. The selection of the reporter strains and bioluminescence measurements were performed as previously described [53]. The raw data, the calculation and statistical evaluations are provided in **Supplemental Dataset 2**.

### Co-IP, LC-MS/MS Analyses and Data Processing

Cells were harvested by centrifugation (5,000 × *g*, 4 °C, 10 min) after 24 h of NirP1-3xFLAG overexpression, resuspended in FLAG buffer (50 mM HEPES-NaOH pH 7; 5 mM MgCl_2_; 25 mM CaCl_2_, 150 mM NaCl; 10% glycerol; 0.1% Tween-20) supplemented with Protease Inhibitor (cOmplete, Roche) and lysed in a Precellys homogenizer (Bertin Technologies). The cleared total cell lysate was subjected to co-IP using ANTI-FLAG M2 affinity agarose gel columns (Sigma). Elution of bound proteins was performed with 500 µL FLAG-peptide (Sigma) solution (100 ng/µL) in TBS buffer at 4 °C. Two independent replicates were prepared. Eluate fractions were measured for protein concentrations with the Coomassie Plus Bradford Assay (Thermo Fisher Scientific), purified by acetone/methanol precipitation [54], lyophilized and stored. For further analysis, samples were resolubilized in denaturation buffer (6 M urea, 2 M thiourea in 100 mM Tris/HCl; pH 8.0) at a final protein concentration of 1 µg/µL. Protein disulfide bonds were reduced with 1 mM dithiothreitol for 45 min and the resulting thiol groups were alkylated with iodoacetamide at a final concentration of 5.5 mM for 45 min. Predigestion of proteins with Lys-C for 3 h was followed by dilution with 4 volumes of 20 mM ammonium bicarbonate buffer, pH 8, and overnight digestion with trypsin, both at protease/protein ratios of 1/100. The resulting peptide solutions were acidified with trifluoroacetic acid to pH 2.5, and an aliquot corresponding to 10 µg peptides was purified by stage tips [55]. For LC‒MS/MS-based protein analysis, 40 ng of ach sample was loaded onto an in-house-made 20 cm column with 70 µm ID, packed with ReproSil-Pur 1.9 µm C18 material (Dr. Maisch, Germany) and separated by RP chromatography on an EASY-nLC 1200 system using 60 min gradients. Eluting peptides were analyzed on an Orbitrap Exploris 480 mass spectrometer (Thermo Fisher Scientific, USA) as described [56].

All raw spectra were processed with MaxQuant software (version 1.6.8.0) at default settings. Peak lists were searched against an in-house modified target-decoy database of *Synechocystis* 6803 with 3,681 protein sequences, including NirP1-3xFLAG. LFQ protein intensities of two independent co-IPs were normalized by corresponding values from whole cell extract measurements, log_2_ transformed and plotted against each other using the Perseus software suite (version 1.6.5.0).

### SDS‒PAGE, Native Gel Electrophoresis and Western Blotting

Proteins were mixed with denaturing or reducing loading dye (5× concentrated: 250 mM Tris-HCl pH 6.8, 25% glycerol, 10% SDS, 500 mM DTT, and 0.05% bromophenol blue G-250). Moreover, 6% ß-mercaptoethanol (v/v) was added fresh to the sample in reducing conditions before heating for 5 min at 95 °C. The protein samples were separated by 12% SDS‒PAGE using the PageRuler^TM^ molecular weight marker (Thermo Fisher). To run samples under nonreducing conditions, a native loading dye (5× concentrated: 30% glycerol, 0.05% bromophenol blue G-250, 150 mM Tris-HCl pH 7.5) was used, and boiling was omitted. To make proteins visible, the gel was stained with *Coomassie* dye. The gel was incubated for 30 min under slight shaking, covered with *Instant Blue Coomassie Protein Stain* (Abcam) and washed overnight with ddH_2_O. Western blots targeting FLAG-tagged proteins were performed using FLAG® M2 monoclonal antibody (Sigma) as described previously [24, 25].

### Nitrate and Nitrite Measurements

Wild type, Δ*nirP1* and *nirP1*-overexpressing cultures were cultivated photoautotrophically in BG11 medium with 5% (v/v) CO_2_ in air as the high CO_2_ condition (HC), harvested and resuspended in new medium and aerated with 0.04% (v/v) CO_2_ in air to start low CO_2_ (LC) conditions. Samples were taken under HC conditions before cells were shifted to LC conditions and then after 0, 2, 4, 6 and 24 h of growth in LC conditions and measured in biological triplicates.

To measure nitrate uptake and nitrite accumulation, cells were washed three times and resuspended in BG11 without NaNO_3_ (BG11-N) lacking CuSO_4_. After incubation for 1 h, only 200 µM NaNO_3_ as the nitrogen source or NaNO_3_ together with 2 mM NH_4_Cl and a final concentration of 2 µM CuSO_4_ were added. Samples were collected at the indicated time points and centrifuged (16,800 × *g*, 5 min, room temperature). Concentrations in the resulting supernatants were determined by IC measurements or with the Griess Reagent Kit (Promega). For IC measurements, samples were mixed with methanol (1:3 (v:v)) and centrifuged three times (16,800 × *g*, 10 min, room temperature), and 10 µl was loaded and separated by a column (DionexTM IonPacTM AS11-HC, Thermo Fischer). Ions were identified, and concentrations were quantified using the area (µS min^-1^) given in the software (Chromeleon® 7). Standards at various concentrations were used for calibration. The raw data, calculations and statistical evaluations are given in **Supplementary Table S3**. Nitrite was measured after conversion into an azo dye [57]. The determination limit was 0.09 µmol L^-1^. The combined standard uncertainty was 4.9%. Samples higher than the measuring range were diluted with ultrapure water.

### Extraction and Metabolite Analysis

Wild type, Δ*nirP1* and *nirP1* overexpressor cultures were cultivated in duplicates without antibiotics and the expression of NirP1 was initiated by the addition of 2 µM CuSO_4_. At the indicated time points, 2 mL samples were collected and snap frozen in liquid nitrogen. For extraction, 630 µL of methanol containing carnitine as an internal standard was added and samples were incubated in a sonic water bath for 5 min. Following phase separation by centrifugation (16,800 × g) for 5 min at room temperature, the upper phase was collected and lyophilized. Absolute metabolite contents were quantified on an HPLC‒MS-8050 system (Shimadzu) as described [58]. The data were further normalized to the OD_750_ measured for each sample and corrected by signals obtained from the wild type. The raw data and estimations as absolute values in ng per mL per OD750 are given in **Supplemental Dataset 4**, with all the statistical evaluations.

Additional detailed materials and methods are provided in **Supplementary Methods**. This includes information on the NirP1-3xFLAG co-IP assay, LC-MS/MS analyses, the extraction and analysis of metabolites and data processing.

## DATA, MATERIALS, AND SOFTWARE AVAILABILITY

The mass spectrometry proteomics data have been deposited to the ProteomeXchange Consortium via the PRIDE [59] partner repository with the dataset identifier PXD041127.

## ACKNOWLEDGMENTS

We are very grateful to Vanessa Krauspe for the previous work on the identification of novel small proteins, Marcus Ziemann for the support in bioinformatic analysis and Annegret Wilde (all Freiburg) for helpful discussions and for access to the spectrophotometer. We thank Ralf Weßbecher (AG Boll, Freiburg) for access and help during the IC measurements.

## CONFLICT OF INTEREST

The authors declare that they have no conflicts of interest.

### Author Contributions

WRH designed the project and secured funding. Construction of mutant and overexpressing strains, characterization of the strains, and co-IP experiments were performed by AW and AK. PS and BM performed and interpreted the MS-based proteomic analyses. Metabolite levels were analyzed by ST and MH. RS performed the shift to LC and the corresponding measurements of nitrite levels in the medium. All other experiments and analyses were performed by AK. AK and WRH wrote the manuscript with input from all authors.

