## Supplement for "Nitrite secretion by cyanobacteria is controlled by the small protein NirP1"

**<u>Content:</u>**

**Supplementary Methods:** **p. 2**

**Supplemental Figures:** **p. 5**

**Supplemental Tables:** **p. 12**

**Overview on Supplemental Datasets:** **p. 15**

**Supplemental References:** **p. 15**

### Supplementary Methods

#### NirP1-3xFLAG co-IP Assay, LC-MS/MS Analyses and Data Processing.

The overexpression of NirP1-3xFLAG was induced in exponentially growing cultures of the *Synechocystis* 6803 strain NirP1oex in the CellDeg system (100 mL at OD ~12) by adding $Cu_2SO_4$ to a final concentration of 2 µM. After another 24 h of cultivation, the cells were harvested by centrifugation (5,000 × *g*, 4 °C, 10 min). Subsequently, the cells were resuspended in FLAG buffer (50 mM HEPES-NaOH pH 7; 5 mM $MgCl_2$; 25 mM $CaCl_2$, 150 mM NaCl; 10% glycerol; 0.1% Tween-20) supplemented with Protease Inhibitor (cOmplete, Roche). For cell lysis, mechanical disruption using a prechilled Precellys homogenizer (Bertin Technologies) was used (3 rounds of shaking at 6,000 rpm (10 sec) followed by a pause (5 sec) per cycle, for a total of 5 cycles, 4 °C). To remove cell debris and glass beads, extracts were centrifuged (1,000 × *g*, 5 min, 4 °C) once, and the supernatant was separated and centrifuged (15,000 × *g*, 30 min, 4 °C) again. The resulting supernatant (i.e., total cell lysate) was subjected to co-IP of NirP1-3xFLAG using ANTI-FLAG M2 affinity agarose gel matrix (Sigma). Therefore, total cell lysates were loaded onto gravity columns packed with 100 µL FLAG agarose and subsequently washed 8 times with 2 mL FLAG buffer. Elution of bound proteins was performed with 500 µL FLAG-peptide (Sigma) solution (100 ng/µL) in TBS buffer (150 mM NaCl in 20 mM Tris/HCl, pH = 7.6) at 4 °C. Two independent replicates were prepared. Protein concentrations of eluate fractions and respective whole cell lysates were measured in the Coomassie Plus Bradford Assay (Thermo Fisher Scientific) through the plate photo reader 1420 multilabel counter (Perkin Elmer) in duplicate, and 500 µL of each was subsequently snap frozen. Proteins in whole cell lysates were purified by acetone/methanol precipitation as described previously [1], and co-IP eluates were lyophilized. All samples were resolubilized in denaturation buffer (6 M urea, 2 M thiourea in 100 mM Tris/HCl; pH 8.0) at a final protein concentration of 1 µg/µL. All subsequent steps were performed at room temperature with shaking at 750 rpm. Protein disulfide bonds were reduced by incubation with dithiothreitol at a final concentration of 1 mM for 45 min, and the resulting Cys thiol groups were alkylated with iodoacetamide at a final concentration of 5.5 mM for 45 min in the dark. Predigestion of proteins with Lys-C for 3 h was followed by dilution with 4 sample volumes of 20 mM ammonium bicarbonate buffer, pH 8, and overnight digestion with trypsin, both at protease/protein ratios of 1/100. The resulting peptide solutions were acidified with trifluoroacetic acid to pH 2.5, and an aliquot corresponding to 10 µg peptides was purified by stage tips [2]. For LC–MS/MS-based protein analysis, 40 ng of ach sample was loaded onto an in-house-made 20 cm column with 70 µm ID, packed with ReproSil-Pur 1.9 µm C18 material (Dr. Maisch, Germany) and separated by RP chromatography on an EASY-nLC 1200 system using 60 min gradients. Eluting peptides were subjected to online coupled electrospray ionization and analyzed in the data-dependent acquisition mode on an Orbitrap Exploris 480 mass spectrometer (Thermo Fisher Scientific, USA) as described previously [3].

All raw spectra were processed with MaxQuant software (version 1.6.8.0) at default settings. The match between the run option and label-free quantification (LFQ) was enabled. Peak lists were searched against an in-house modified target-decoy database of *Synechocystis* 6803 with 3,681 protein sequences, including the NirP1-3xFLAG fusion protein sequence. False discovery rates were limited to 1% at the peptide and protein levels. LFQ protein intensities of two independent co-IPs were normalized by corresponding values from whole cell extract measurements, $\log_2$ transformed and plotted against each other using the Perseus software suite (version 1.6.5.0).

**Extraction and Metabolite Analysis.**

Wild type, Δ*nirP1* and *nirP1* overexpressor cultures were set in duplicates to an OD of 0.5 and grown for 24 h without antibiotics before cells were pelleted by centrifugation (3,210 × g, 10 min, room temperature), washed three times in copper-free BG11 medium and induction of NirP1 was started by adding 2 µM $CuSO_4$ after 1 h.

After 0, 1, 3, 12 and 24 h of incubation, 2 mL of each culture was collected by centrifugation (16,800 × g, 5 min, room temperature) and immersed in liquid nitrogen. For extraction, 630 µL of methanol containing carnitine (internal standard, 1 mg per extraction) was added, mixed for 1 min, and incubated for 5 min in a sonic water bath. After another 15 min of shaking at room temperature, 400 µL of chloroform was added, and the sample was incubated at 37 °C for 5 min. Next, 800 µL of ROTISOLV LC–MS grade $H_2O$ (Roth) was added, and the sample was mixed thoroughly. Precipitation was enabled by storage at 22 °C for at least 2 h, and phase separation was subsequently achieved by centrifugation for 5 min at room temperature (16,800 × g). The upper polar phase was collected and dried in a speed vac for 30 min followed by lyophilization overnight.

Absolute metabolite contents were quantified on a high-performance liquid chromatography–mass spectrometer LC–MS-8050 system (Shimadzu) as described previously [4]. Dried samples were dissolved in 200 µL LC–MS grade water and filtered through 0.2-mm filters (Omnifix-F, Braun, Germany), and 5 µL of the cleared supernatant was separated on a pentafluorophenylpropyl column (Supelco Discovery HS FS, 3 mm, 150 3 2.1 mm).

The compounds were identified and quantified using the multiple reaction monitoring (MRM) values given in the LC–MS/MS method package and the LabSolutions software package (Shimadzu). Authentic standard substances (Merck) at various concentrations were used for calibration, and peak areas were normalized to signals of the internal standard (carnitine). The data were further normalized to the $OD_{750}$ measured for each sample and corrected by signals obtained from the wild type. The raw data and estimations as absolute values in ng per mL per OD750 are given in **Supplemental Dataset 4**, with all the statistical evaluations.

### Supplemental Figures

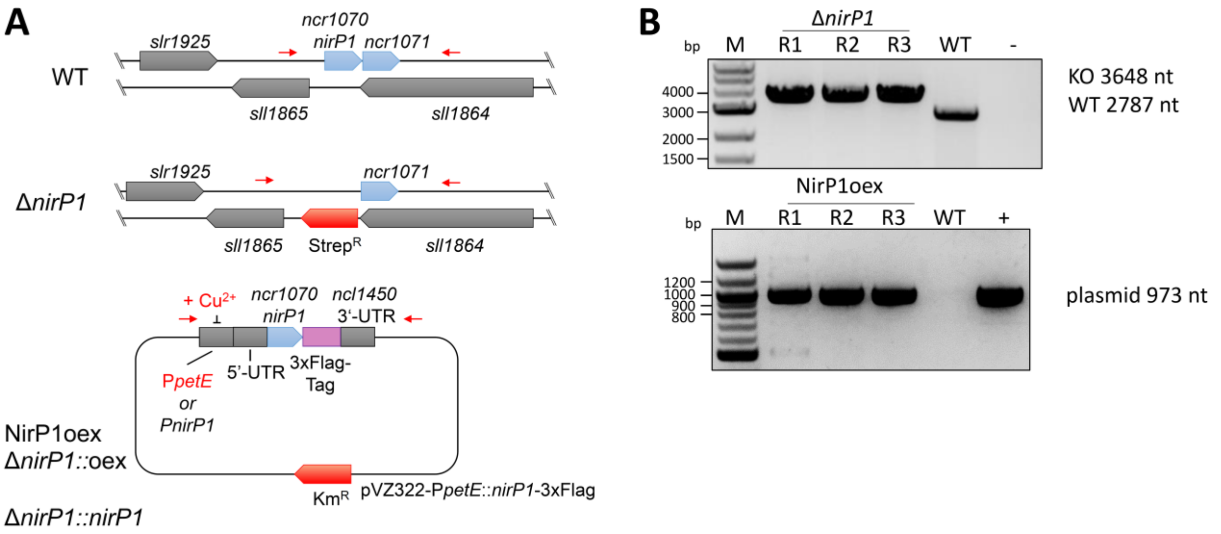


**Fig. S1. Generation of Δ*nirP1* and *nirP1*-overexpressing strains. A** The *nirP1* locus in the wild type and in the Δ*nirP1* deletion strain, as well as of a pVZ322 plasmid derivative harboring a *nirP1* gene copy under the control of the $Cu^{2+}$-inducible $P_{petE}$ promoter in the *nirP1* overexpression strains (NirP1oex and Δ*nirP1::*oex) and the complementation strain (Δ*nirP1:: nirP1*) under the control of its native promoter. The *nirP1* gene was previously assigned to the transcriptional unit (TU) 2296 [5] extending from position 2215954 to 2217039 on the forward strand of the *Synechocystis* 6803 chromosome (GenBank accession no. NC_000911). The first segment of TU2296 has also been annotated as ncr1070 due to its sometimes-divergent accumulation compared to the second half, overlapping gene *sll1864* on the reverse strand. In the Δ*nirP1* strain, the gene was replaced by a streptomycin resistance cassette ($Strep^R$) via homologous recombination. The plasmid enabling ectopic *nirP1* expression was introduced into *Synechocystis* wild type and Δ*nirP1* strains. The red arrows indicate primer binding sites used to verify the mutants. **B** PCR verification of the genotypes of independently obtained mutant strains. Three clones were tested each, using primers P_AK9/P_AK10 (for Δ*nirP1*) or P_AK16/ P_AK17 (overexpression and complementation strains NirP1oex, Δ*nirP1*::oex and Δ*nirP1*::*nirP1*; primer sequences in **Table S1**). The PCR of NirP1oex representative for all pVZ322-containing strains is shown. Abbreviations: M, marker; bp, base pairs; c., clone; -, negative control (water control); +, positive control (purified plasmid as template). This figure is an extension to **Fig. 1** and illustrates technical aspects.

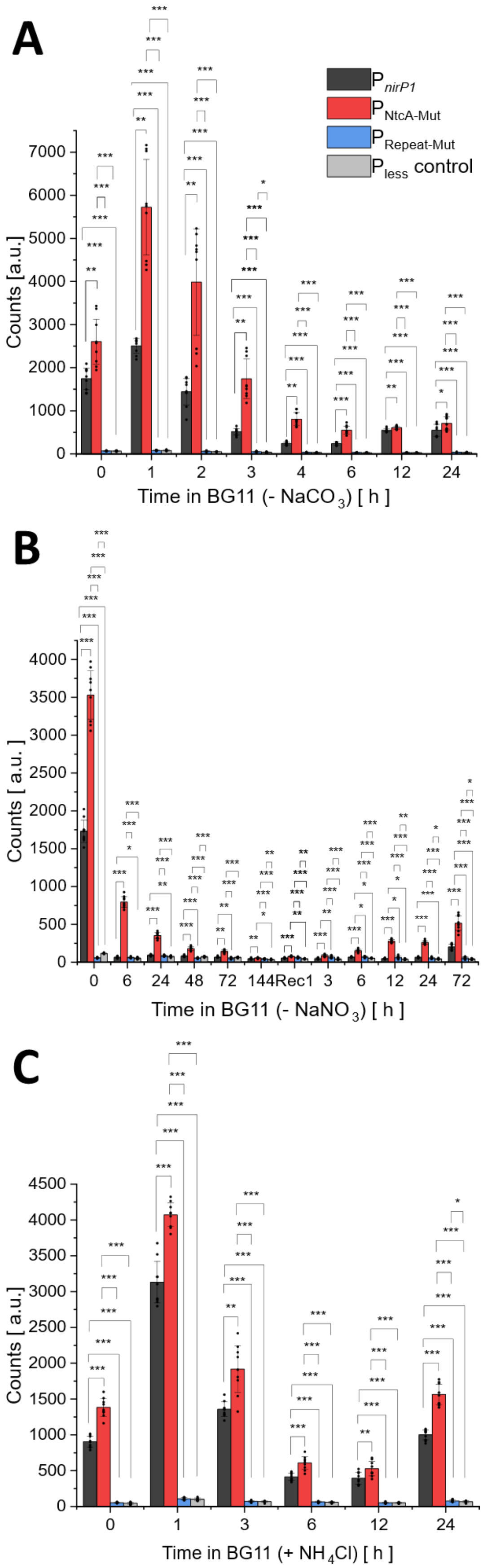


**Fig. S2. NirP1 expression is mediated through NtcA and a $C_i$-sensitive promoter.**

**A** Bioluminescence of the *Synechocystis* 6803 reporter strains. Initially, cells were grown under HC for 2 h and transferred to BG11 medium without $CO_2$ source to induce LC conditions for 24 h. To improve the signal of bioluminescence mediated through the promoters, 2 µL decanal were added to the measurements. Bioluminescence data are presented as the means ± SD of three independent measurements for three biological replicates each (resulting from three independent transformants). In each experiment, a strain carrying a promoterless (pless) *luxAB* was used as a negative control (in each case measured in three independent cultures, n=3). **B** Cells were cultivated in standard conditions and subsequently transferred to $NO_3$-free BG11 medium to start nitrogen deprivation. After 7 days (time point 144 h) of nitrogen starvation, bleached cells were transferred back to standard BG11 medium (indicated as Rec) to start the recovery. **C** Cells were grown in standard conditions $T_0$ and were further grown with additional with 10 mM $NH_4Cl$ for 24 h. Significance was calculated with a two-sample *t* test with unequal variance (Welch's *t* test; *, $P < 0.05$; **, $P < 0.01$; ***, $P < 0.001$) between the strains at corresponding time points. Further details of statistical analysis are given in **Supplemental Dataset 2**.

These data represent an extension to the results shown in **Fig. 2**.

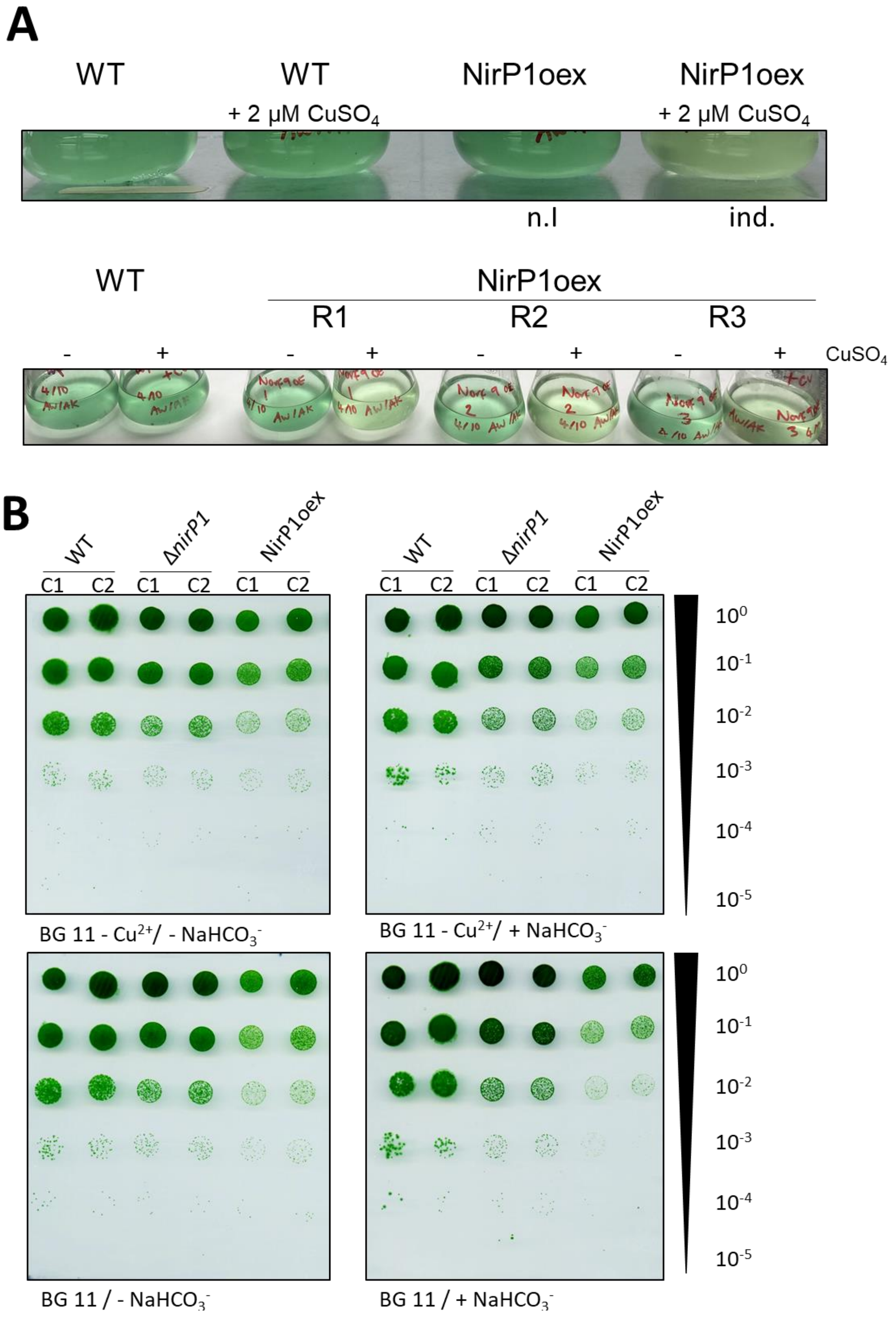


**Fig. S3. Phenotypical differences between *nirP1* mutants and the wild type. A** Pigmentation phenotype of wild type and *nirP1* overexpressor (NirP1oex) without and with 2 µM $Cu_2SO_4$ to trigger the expression of *nirP1* for 24 h from the $P_{petE}$ promoter.

Lower panel: Repetition with three biological replicates of NirP1oex (R1 to R3). WT was used as negative control. **B** Drop dilution assay on solid medium comparing duplicate samples of the *Synechocystis* wild type (WT), the *nirP1* deletion mutant Δ*nirP1* and overexpression strain NirP1oex under continuous light in LC (without $NaHCO_3$) and HC (with 10 mM $NaHCO_3$) conditions. Strains were pre-cultivated in liquid BG11 medium without $Cu_2SO_4$ and without $NaHCO_3$ under constant light, and the indicated different dilutions were spotted on agar plates. For ectopic overexpression of NirP1, standard BG11 medium containing 0.3 μM $Cu_2SO_4$ was used in the plates. Plates were analyzed after incubation for 5 days.

These data represent an extension to the results shown in **Fig. 3**.

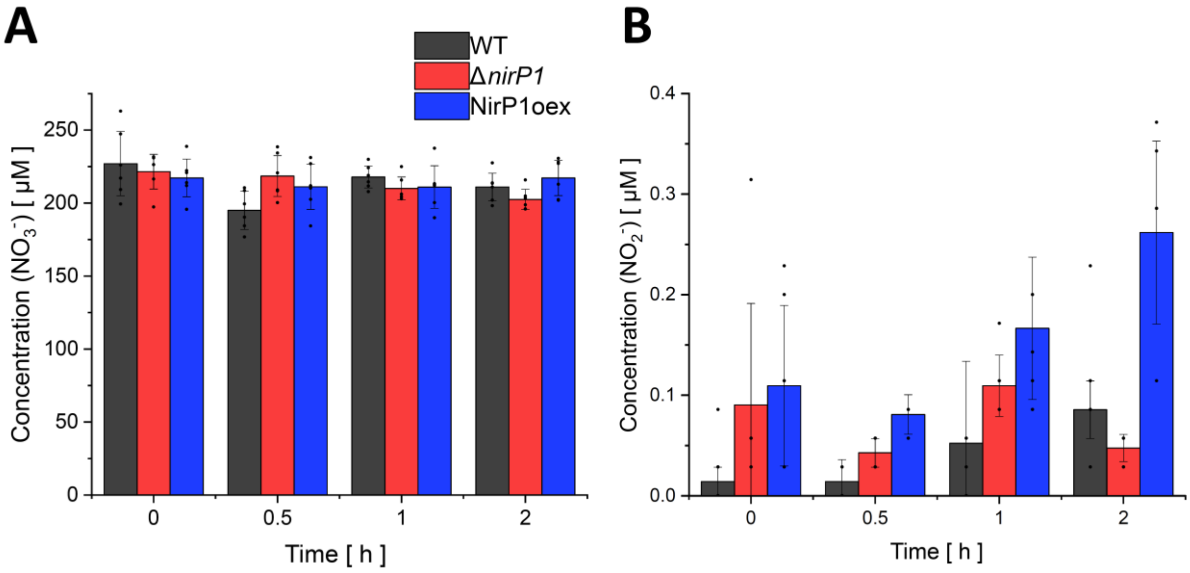


**Fig. S4. Uptake of $NO_3$ and secretion of $NO_2$ in the presence of ammonia**. **A** Uptake of $NO_3$. **B** Secretion of $NO_2$. The strains were grown in BG11, washed with nitrogen-free BG11 (BG11 –N), set to OD = 1 and resuspended in BG11 containing 200 µM nitrate and 2 mM ammonium. Concentration of nitrate and nitrite was measured in the supernatant after centrifugation of the cell suspensions. The data are presented as the means ± SD of two independent measurements for three biological replicates each (resulting from independent transformants). Data details and statistical analysis of nitrate consumption and the nitrite excretion assay are given in **Supplemental Dataset 3**.

This is a control experiment to **Fig. 6**.

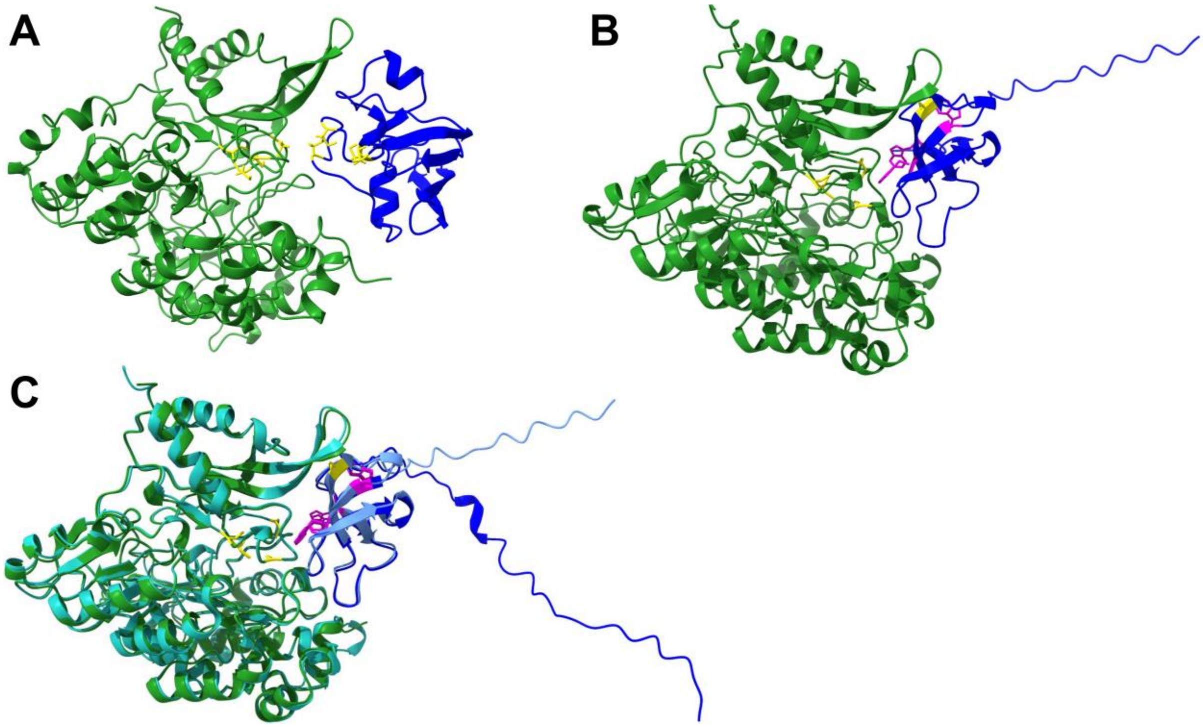


**Fig. S5. Predicted NiR interactions with Fd1 in *Synechocystis* 6803 and with NirP1 in *Synechococcus* 7942. A** Predicted interaction of the 97 AA sequence WP_190595418.1 for Fd1 (*ssl0020, petF*) and the 502 AA sequence WP_010873675.1 for NiR from *Synechocystis* 6803 using AlphaFold [6, 7]. Both structures were modeled from the first to the last AA. The residues of the iron-sulfur clusters in each protein are highlighted in yellow. **B** Predicted interaction between the *Synechococcus elongatus* sp. PCC 7942 NiR-NirP1 homologs generated by AlphaFold using the 75 AA sequence WP_011244559.1 for NirP1 and the 512 AA sequence WP_011242624.1 for NiR. The residues of the iron-sulfur clusters in NiR are highlighted in yellow and the conserved AA in the NirP1 homologue as in **Fig. 5E** for the *Synechocystis* 6803 NirP1. (*C*) Overlay of the predicted interactions between NirP1 and NiR from *Synechocystis* 6803 *(*NiR in green and NirP1 in blue*)* and *Synechococcus (*NiR in black and NirP1 in orange*)* showing similar arrangements in the two different organisms.

This Figure is an extension to **Fig. 5E.**

### Supplemental Tables

**Table S1.** Desoxyoligonucleotide primers used in this study (abbreviations: AQ, AQUA cloning [8]).

| Name | Sequence (5‘ to 3‘) | Description | Purpose |
|---|---|---|---|
| P_AK1 | CACAGCTTGTACTTAGCTTTGAGAATGG | AQ of *nirP1* KO construct to create flank1 (F1) with overlap to puC19 plasmid backbone | Generation of the *nirP1* knock out strain with pUC19 plasmid |
| P_AK2 | TACGATAATATTACAAAATGATAGGGATAATAATTTG | AQ of *nirP1* KO to create flank1 with overlap to the streptomycin resistance cassette (Strep$^{R}$) | |
| P_AK3 | CATTTTGTAATATTATCGTAGTTGCTCTCAG | AQ of *nirP1* KO to create the Strep$^{R}$ cassette with overlap to flank 1 | |
| P_AK4 | CTTGGGCAGCGTACAGAGTGATGTCAAC | AQ of *nirP1* KO to create the Strep$^{R}$ cassette with overlap to flank 2 | |
| P_AK5 | CACTCTGTACGCTGCCCAAGCTAGAATC | AQ of *nirP1* KO to create flank 2 (F2) with overlap to the Strep$^{R}$ cassette | |
| P_AK6 | CCGCTTACAGCCCCCAATCCCATCACTG | AQ of *nirP1* KO to create flank 2 with overlap to the Strep$^{R}$ cassette | |
| P_AK7 | GGATTGGGGGCTGTAAGCGGATGCCGGG | AQ of *nirP1* KO to create the backbone of puC19 with overlay to flank 2 | |
| P_AK8 | AAAGCTAAGTACAAGCTGTGACCGTCTC | AQ of NirP1 KO to create the backbone of puC19 with overlay to flank1 | |
| P_AK9 | CATGTCCCCTCGAATTTC | Segregation check for *nirP1* deletion | |
| P_AK10 | GTGCACATTGCCCTTAAG | Segregation check for *nirP1* deletion | |
| P_AK11 | GACATTAACCTATAAAAATAGGCG | Sequencing the flank 1 of NirP1 KO construct in puc19 | |
| P_AK12 | GGCAGGTCAAAGATAGTC | Sequencing the Strep$^{R}$ cassette of NirP1 KO construct in puc19 | |
| P_AK13 | GATCAAAGAGTTCCTCCG | Sequencing the Strep$^{R}$ cassette of NirP1 KO construct in puc19 | |
| P_AK14 | GCGGAAAATAAACACAGTGG | Sequencing flank 2 of NirP1 KO construct in puc19 | |
| P_AK15 | GTAATAAAATGCGCCCTTC | Sequencing the last part of Flank 2 of NirP1 KO construct in puc19 | |
| P_AK16 | CTGCCCGGATTACAGATC | Sequencing the *nirP1* constructs on the pVZ322 plasmid | Construction of pVZ322::$P_{nirP1/petE}$::nirP1::3xFLAG for ectopic expression in *Synechocystis* |
| P_AK17 | GTAATACCATGAAAAATACCATGCTCAG | Sequencing the *nirP1* constructs on the pVZ322 plasmid | |
| P_AK18 | GGATTACAGATCCTCTAGAGACTTATCGTGTTTGTCAAG | Fragment for the native promotor of NirP1 with the overlap to the pVZ322 plasmid | |

| | | | |
|---|---|---|---|
| P_AK19 | AGGTTCATTTGTGGCGACATAAAGTCCTCTCCATTTTAC | Fragment for the native promotor of NirP1 with the overlap to the pVZ322 plasmid | |
| P_AK20 | ATGTCGCCACAAATGAACC | Primer for the ORF of NirP1 at the translational start site | |
| P_AK21 | CCATCATGATCTTTATAATCACAATAAAAATCACTGCGGTC | Primer for the ORF of NirP1 | |
| P_AK22 | TTACAGATCCTCTAGAGTCGACCTGGGCCTACTGGGCTATTC | Fragment for the *petE* promotor of NirP1 with the overlap to the pVZ322 plasmid | |
| P_AK23 | GGTTCATTTGTGGCGACATACTTCTTGGCGATTGTATC | Fragment for the petE promotor of NirP1 with the overlap to the pVZ322 plasmid | |
| P_AK24 | GATTATAAAGATCATGATGG | 3xFlagTag and 3'UTR for taggig small proteins | |
| P_AK25 | GATGTATGCTCTTCTGCTCCTGCAGTAATAAAAAACGCCCGGCGGCAACCGAGCGAATAATTCCCAACGAAGGCAAGC | 3xFlagTag and 3'UTR for taggig small proteins | |
| P_AK26 | GTGCAGGTCGACTCTACCGGACTTATCGTGTTTGTCAAGATTTG | Promotor site of NirP1 (region from -70 to + 30) | pILA vectors with *nirP1* promotor site and variants for biolumi-nescence assays |
| P_AK27 | TCATCTAATGCTAAGGCCGGAAGTCAAAGGAATAATTTCTGTATCC | Promotor site of NirP1 (region from -70 to + 30) | |
| P_AK28 | TCATCTAATGCTAAGGCCGGAAGTCAAAGGAATAATTTCTGATTCCATATTTAGAAAATG | Promotor site of NirP1 with mutation in the NtcA binding site (region from -70 to + 30) | |
| P_AK29 | GTGCAGGTCGACTCTACCGGACTTATCGTGTTTGTCCCGAGGGCTTAATTTTG | Promotor site of NirP1 with mutation in the tandem repeat (region from -70 to + 30) | |

**Table S2.** Plasmids used in this study.

| Plasmid name | Description | Reference |
|---|---|---|
| pUC19 | High-copy cloning vector used for replication in *E. coli*, $Amp^R$ | [9] |
| pUC19::*nirP1*_KO | Plasmid used for homologous recombination of $kan^R$ cassette into the *nirP1* locus using up- and downstream genomic regions flank 1 and flank 2 as flanking sites to create the KO strain Δ*nirP1* | This study |
| pVZ322::$P_{nirP1}$::*nirP1*::3×FLAG | Self-replicating, conjugative plasmid for native *nirP1* expression in *Synechocystis* 6803 driven by $P_{nirP1}$ promoter, $Km^R$ & $Gen^R$ | This study |
| pVZ322::$P_{petE}$::*nirP1*::3×FLAG | Self-replicating, conjugative plasmid for copper inducible *nirP1* expression from $P_{petE}$ promotor in *Synechocystis* 6803, $Km^R$ & $Gen^R$ used to create the expression strains NirP1oex and Δ*nirP1*::*nirP1* | This study |
| pilA (FseI, AgeI) | Plasmid used for homologous recombination of promotor regions fused to *luxAB* genes; used for bioluminescence assay | [10, 11] |
| pILA::$P_{nirP1}$::*luxAB* | Plasmid to fuse the *nirP1* promotor region to *luxAB* genes for bioluminescence assay | This study |
| pILA::$P_{NtcA\text{-}Mut}$::luxAB | Plasmid to fuse the *nirP1* promotor region with mutation in the NtcA binding site to *luxAB* for bioluminescence assay | This study |
| pILA::$P_{Repeat\text{-}Mut}$::luxAB | Plasmid to fuse the *nirP1* promotor region with mutations in the upstream binding motif to *luxAB* for bioluminescence assay | This study |

**Table S3. List of identified proteins from NirP1-3xFLAG Co-IP and corresponding LFQ intensities.** See separate Excel file "**Supporting Tables S3-5.xlsx**".

**Table S4. List of identified peptides of NirP1-3xFlag (Norf9-3xFlag).** See separate Excel file "**Supporting Tables S3-5.xlsx**".

**Table S5. List of identified peptides of NiR.** See separate Excel file "**Supporting Tables S3-5.xlsx**".

### Overview on Supplemental Datasets

**Supplemental Dataset 1.** Sequences of 485 detected homologs of NirP1. See separate Excel file “Supplemental Dataset1_2_3.xlsx”.

**Supplemental Dataset 2**. Measurements and statistical analysis of the promoter bioluminescence assay. See separate Excel file “Supplemental Dataset1_2_3.xlsx”.

**Supplemental Dataset 3**. Measurements and statistical analysis of the nitrate uptake and nitrite accumulation experiment. See separate Excel file “Supplemental Dataset1_2_3.xlsx”.

**Supplemental Dataset 4**. Measurements and statistical analysis of extracted metabolites. See separate Excel file “Supplemental Dataset4.xlsx”..
